# Immunogenicity of a public neoantigen derived from mutated *PIK3CA*

**DOI:** 10.1101/2021.04.08.439061

**Authors:** Smita S. Chandran, Jiaqi Ma, Martin G. Klatt, Friederike Dundar, Chaitanya Bandlamudi, Pedram Razavi, Hannah Y. Wen, Britta Weigelt, Paul Zumbo, Si Ning Fu, Lauren B. Banks, Watchain D. Bestman, Alexander Drilon, Doron Betel, David A. Scheinberg, Brian M. Baker, Christopher A. Klebanoff

**Author notes:** Email correspondence to C.A.K. or S.S.C.; Address: Memorial Sloan Kettering Cancer Center, 417 E 68th St, Office #763. New York, NY, 10065.

## Abstract

Public neoantigens (NeoAgs) represent an elite class of shared cancer-specific epitopes derived from recurrent mutations in driver genes that are restricted by prevalent HLA alleles. Here, we report on a high-throughput platform combining single-cell transcriptomic and T cell receptor (TCR) sequencing to establish whether mutant (Mut) *PIK3CA*, among the most common genomically altered driver oncogenes, generates an immunogenic public NeoAg. Using this method, we developed a library of TCRs that recognize an endogenously processed neoepitope containing a common *PIK3CA* hotspot mutation that is restricted by HLA-A*03:01. Mechanistically, immunogenicity to this public NeoAg arises primarily from enhanced stability of the neopeptide/HLA complex caused by a preferred HLA anchor substitution. Structural studies indicated that the HLA-bound neopeptide presents a relatively “featureless” surface dominated by the peptide’s backbone. To overcome the challenge of binding such an epitope with high specificity and affinity, we discovered that a lead TCR clinical candidate engages the neopeptide through an extended interface aided by an unusually long β-chain complementarity-determining region 3 (CDR3β) loop. In a pan-cancer cohort of patients with diverse malignancies that express the *PIK3CA* public NeoAg, we observed spontaneous immunogenicity, NeoAg clonal conservation, and in a limited number of cases, evidence of targeted immune escape. Together, these results establish the immunogenic potential of Mut *PIK3CA*, creating a framework for off-the-shelf immunotherapies targeting this public NeoAg.

## INTRODUCTION

Neoantigens (NeoAgs), human leukocyte antigen (HLA)-bound peptides resulting from the protein products of non-synonymous somatic mutations (NSSMs), are a major class of human cancer regression antigens^1,2^. In >99% of cases, NeoAgs are exclusive to individual patients because they result from random passenger mutations that do not contribute to cancer cell fitness^3^. Patient-specific, or “private”, NeoAgs represent a significant challenge to therapeutically target in a time and cost-efficient manner because they must be customized^4^. Further, selective loss of tumor clones that express immunogenic private NeoAgs can increase intratumoral heterogeneity (ITH), a major mechanism of cancer immunotherapy resistance^5–8^. In contrast, NeoAgs resulting from recurrent mutations in driver genes would generally be clonally conserved because these alterations directly contribute to cancer initiation, growth, and survival^9^. If a NeoAg resulting from a driver hotspot mutation is also restricted by a prevalent HLA allele, this elite subset of NeoAgs would be shared among cancer patients, creating a “public” NeoAg^10^.

The first human public NeoAg was described 25 years ago resulting from an HLA-A*02:01-restricted epitope derived from a recurrent cyclin-dependent kinase 4 (*CDK4*) mutation^11^. Since that time, only a limited number of additional public NeoAg examples have been reported^12–18^; collectively, these are associated with <3% of the 299 driver genes contained in The Cancer Genome Atlas (TCGA) consensus list^19^. Critically, none have previously been reported arising from mutant (Mut) *PIK3CA* (the gene encoding the alpha subunit of phosphatidylinositol 3-kinase, PI3Kα), among the most common genetically-altered driver oncogenes in humans^19–21^. Small molecule PI3K inhibitors cause cancer regression in patients, validating PI3Kα as a therapeutic target^22^. However, these inhibitors are not curative and are associated with significant on-target but off-tumor toxicities. Novel immunotherapies that specifically target Mut PI3Kα but spare heathy tissues could therefore have broad utility for many cancer patients.

Herein, we report the development and application of a high-throughput platform that combines single-cell transcriptomic and paired α/β T cell receptor (TCR) sequencing to test whether Mut *PIK3CA* generates an immunogenic public NeoAg. Using this method, we developed a library of TCRs that recognize an endogenously processed neoepitope resulting from a common *PIK3CA* hotspot mutation. Each TCR library member is restricted by HLA-A*03:01, among the most prevalent HLA alleles in North America and Europe^23^, thereby establishing a *PIK3CA* public NeoAg. Structural and biophysical measurements revealed that the mechanism of immunogenicity for this shared NeoAg arises primarily from enhanced stability of the neopeptide/HLA complex caused by an optimal substitution at a primary HLA anchor position. These studies additionally revealed that the HLA-bound neopeptide presents a relatively “featureless” surface dominated by the peptide backbone. To overcome the challenge of engaging such a neopeptide with a high specificity and relatively high affinity, we discovered that a lead TCR clinical candidate engages the HLA complex in a unique extended fashion by incorporating an unusually long hypervariable β-chain complementarity-determining region 3 (CDR3β) loop. Finally, by analyzing a cohort of cancer patients who express the *PIK3CA* public NeoAg, we demonstrate evidence of spontaneous immunogenicity, NeoAg clonal conservation, and in a limited number of cases, immune escape. Together, these results establish the immunogenic potential of a shared neoantigen derived from Mut *PIK3CA*, creating a framework for future off-the-shelf immunotherapies targeting this public NeoAg.

## RESULTS

### A high-throughput single-cell Mut PIK3CA-specific TCR discovery platform

We developed a modular, high-throughput discovery platform to simultaneously test whether Mut PI3Kα is immunogenic and to retrieve paired α/β TCR gene sequences that confer specificity to this NeoAg (**Fig. 1A**). The platform proceeds in four steps. First, naive T cells (T_N_) from a healthy donor (HD) undergo *in vitro* sensitization (IVS) with autologous monocyte-derived dendritic cells (moDCs) electroporated with mRNA encoding the Mut driver gene. This approach was designed to empirically test whether Mut PI3Kα is capable of endogenous proteolytic processing and HLA presentation. It is agnostic of *in silico* peptide/HLA binding algorithms and does not require *a priori* knowledge of a minimal epitope. Second, matched aliquots from sensitized wells are re-stimulated with moDCs expressing the Mut or WT version of the driver. Transcripts for *IFNG*, the gene encoding interferon-γ, are quantified by qPCR and a regression analysis is performed as a screen to identify “hit” wells containing Mut-specific T cells. Third, T cells from “hit” wells are again acutely re-stimulated with moDCs to identify reactive clonotypes and retrieve their TCR gene sequences. This step, which we have termed **<u>S</u>**timulation **I**nduced **<u>F</u>**unctional **<u>T</u>**CR **<u>seq</u>**uencing (SIFT-seq), is accomplished using combined single-cell (sc) TCR V(D)J and RNA sequencing. Finally, TCR clonotypes that selectively express T cell activation transcripts in response to a Mut variant of the driver but not its WT counterpart are reconstructed for functional validation.

**Fig. 1:**
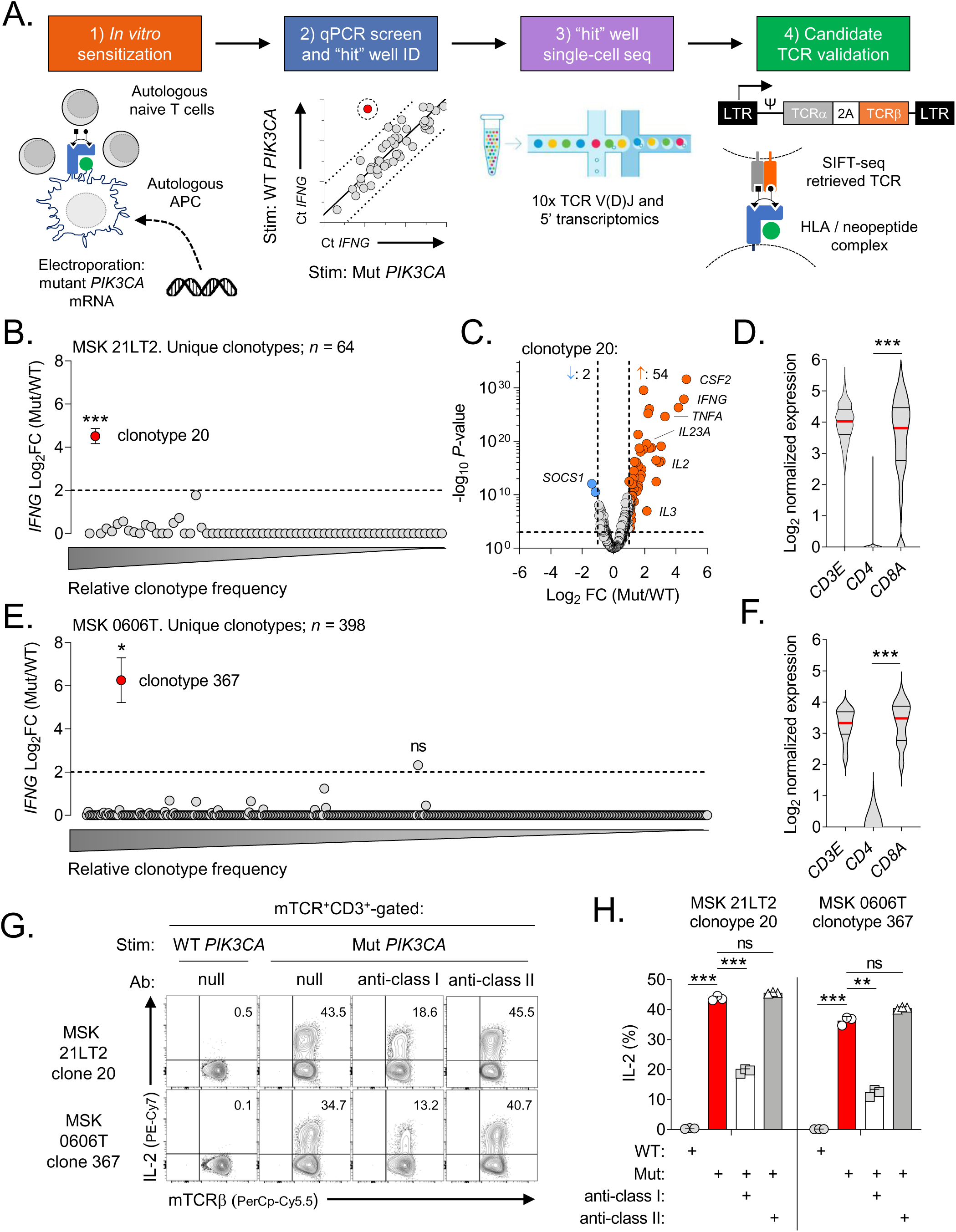
Development and validation of the SIFT-seq discovery platform for Mut *PIK3CA*-specific TCRs. (**A**) Schematic overview of the SIFT-seq TCR discovery platform. **(B)** Log_2_ fold-change (FC) ratio of *IFNG* transcripts from *n*=64 unique TCR clonotypes identified using single-cell RNA and V(D)J TCR sequencing from screen positive “hit” well MSK 21LT2. Matched aliquots of sensitized T cells from healthy donor (HD) 1 were stimulated with *PIK3CA* (H1047L) (Mut) or wildtype *PIK3CA* (WT) prior to single-cell sequencing. The mean *IFNG* ratio for all evaluable TCR clonotypes is shown. Statistical analyses were performed for all clonotypes with a minimal ratio ≥2 (dashed line). The X-axis indicates the relative frequencies of individual detected TCR clonotypes. (**C**) Volcano plot displaying global transcriptomic changes for MSK 21LT2 clonotype 20 following stimulation with Mut versus WT *PIK3CA*. Vertical and horizontal dashed lines indicate thresholds for gene expression FC and statistical significance, respectively. Orange and blue dots represent significantly up and down-regulated genes following Mut *PIK3CA* stimulation, respectively. (**D**) Violin plots depicting transcript levels for the lineage markers *CD3E*, *CD4* and *CD8A* from MSK 21LT2 clonotype 20. (**E**) Log_2_ fold-change (FC) ratio of *IFNG* transcripts from *n*=398 unique TCR clonotypes identified using single-cell sequencing from screen positive “hit” well MSK 06006T derived from HD2. (**F**) Violin plots depicting lineage marker transcript expression for MSK 0606T clonotype 367. (**G**) Representative FACS plots and (**H**) summary bar graph (*n*=3 replicates per condition) displaying the frequency of intra-cellular IL-2 production in open-repertoire T cells retrovirally transduced with SIFT-seq retrieved TCR candidates. The reconstructed TCR expresses a murine constant chain (mTCR), enabling detection with an anti-mTCR-specific antibody. Transduced T cells (live^+^mTCR^+^CD3^+^) were co-cultured with autologous monocyte derived DCs electroporated with mRNA encoding Mut or WT *PIK3CA* in the absence or presence of pan-HLA class I or class II blocking antibodies. (**B, E, H**) Symbols and bar graphs displayed as mean ± SEM. * *P* ≤ 0.05; ** = *P* ≤ 0.01; *** = *P* ≤ 0.001. ns = not significant. Two-sided Student’s t-test with Bonferroni correction.

The majority of genomic alterations in *PIK3CA* result from NSSMs occurring at one of three hotspots^24^. Across cancer types, the most common hotspot affects the H1047 residue of the protein’s kinase domain. Additional hotspots alter PI3Kα’s helical domain at residues E542 and E545. Healthy donor-derived T_N_ underwent IVS by moDCs individually transfected with one of the four most frequent *PIK3CA* hotspot mutations (H1047L/R, E542K, and E545K). The qPCR screen identified a single well (21LT2) containing T cells that preferentially upregulated *IFNG* in response to *PIK3CA* (H1047L) (henceforth Mut *PIK3CA*) relative to WT *PIK3CA*. Matched aliquots of T cells from well 21LT2 were subsequently subjected to SIFT-seq. To retrieve candidate TCR gene sequences of Mut *PIK3CA*-specific T cells, we plotted the ratio of *IFNG* produced under Mut and WT stimulation conditions for each detected clonotype (**Fig. 1B**). Among 64 unique clonotypes, only one (clonotype 20) significantly upregulated *IFNG* in response to Mut *PIK3CA*. Global transcriptomic analysis of clonotype 20 revealed additional Mut-specific changes in the expression of genes previously associated with acute TCR activation^25^, including *TNFA*, *IL2*, and *CSF2* (**Fig. 1C**). By contrast, all other clonotypes retained a quiescent transcriptomic profile (**Extended Data Fig. 1A** and data not shown). Lineage marker assessment of clonotype 20 revealed expression of *CD3E*, as expected, and *CD8A* but minimal to no *CD4* (**Fig. 1D**). These data suggested that clonotype 20 was likely HLA class I (HLA-I)-restricted.

To assess the platform’s capacity to reproducibly identify Mut *PIK3CA*-specific TCR clonotypes, we performed an independent run using peripheral blood mononuclear cells (PBMCs) from a second HD (HD2). A qPCR screen positive well (0606T) was identified from HD2 and subjected to SIFT-seq analysis. Among 398 evaluable clonotypes, only a single T cell clone (clonotype 367) selectively upregulated *IFNG* (**Fig. 1E**) and other TCR activation-induced genes (**Extended Data Fig. 1B**) in response to Mut. Similar to clonotype 20 from HD1, clonotype 367 expressed high levels of the *CD3E* and *CD8A* lineage markers but not *CD4* (**Fig. 1F**) suggesting this TCR was also HLA-I restricted. In both runs, the reactive TCR candidate was not the most frequent clonotype within screen positive wells (**Extended Data Fig. 1A** and **C**).

SIFT-seq retrieved TCR gene sequences were subsequently cloned into retroviral expression vectors to establish whether they confer Mut *PIK3CA* reactivity when transduced into open-repertoire T cells. Transduced T cells were subsequently co-cultured with autologous moDCs transfected with Mut or WT *PIK3CA*. Expression of TCRs derived from 21LT2 clonotype 20 and 0606T clonotype 367 resulted in cytokine production exclusively to Mut but not WT *PIK3CA* (**Fig. 1G**), validating the SIFT-seq approach. As controls, we synthesized the dominant clonotypes from both screen positive wells and confirmed these TCR sequences do not confer Mut *PIK3CA* reactivity (**Extended Data Fig. 1D, E** and data not shown). Finally, to determine whether retrieved Mut *PIK3CA*-specific TCRs are HLA-I or -II restricted, we tested their function in the presence or absence of HLA class-specific blocking antibodies. We discovered that the function of both TCRs was attenuated by an HLA-I but not -II blocking antibody (**Fig. 1G and H**). These data indicated that the sc transcriptomic assessment of T cell lineage markers correctly anticipated the HLA-I restriction of the *PIK3CA* Mut-specific clonotypes. Taken together, we conclude that the SIFT-seq platform can reproducibly identify TCR gene sequences that confer specificity to a recurrent *PIK3CA* hotspot mutation.

### HLA restriction and functional characterization of PIK3CA public NeoAg-specific TCRs

We next sought to resolve the specific HLA-I allele(s) required for presentation of a Mut *PIK3CA*-derived epitope to the SIFT-seq retrieved TCRs. High-resolution HLA-I genotyping of HD1 revealed the subject was fully heterozygous at the *HLA-A*, -*B*, and -*C* loci (**Fig 2A**). Among North American and European populations, the frequency of the individual HLA-I alleles expressed by HD1 vary from being highly prevalent to rare^23^. To deconvolute which allele is required for TCR recognition, we generated a panel of mono-allelic target cells expressing individual HLA-I molecules in combination with either Mut or WT *PIK3CA*. Co-culture of this panel with T cells transduced with the 21LT2 clonotype 20 TCR revealed cytokine production exclusively in the context of Mut *PIK3CA* and HLA-A*03:01. This HLA-I allele is among the most common in North America and Europe, occurring in 20%-28% of individuals; the allele is also represented in other populations globally^23^. Cytokine production was not observed in response to target cells expressing either HLA-A*03:01 and WT *PIK3CA* or an alternative HLA-A allele and Mut *PIK3CA*. Using a similar HLA deconvolution method, we resolved that 0606T clonotype 367 is also restricted by HLA-A*03:01 (data not shown). Based on these data, we performed SIFT-seq analysis on additional HLA-A*03:01^+^ HDs and successfully retrieved two additional TCRs that recognized the same Mut *PIK3CA*/HLA-I combination. The resulting Mut *PIK3CA*-specific TCR library (TCRs 1-4) was composed of unique gene sequences derived from diverse V segments and distinct CDR3 lengths (**Extended Data Fig. 2A**). Together, these data establish that the protein product of a *PIK3CA* hotspot mutation presented in the context of a prevalent HLA-I allele can reproducibly elicit a T cell response, creating an immunogenic *PIK3CA* public NeoAg.

**Fig. 2:**
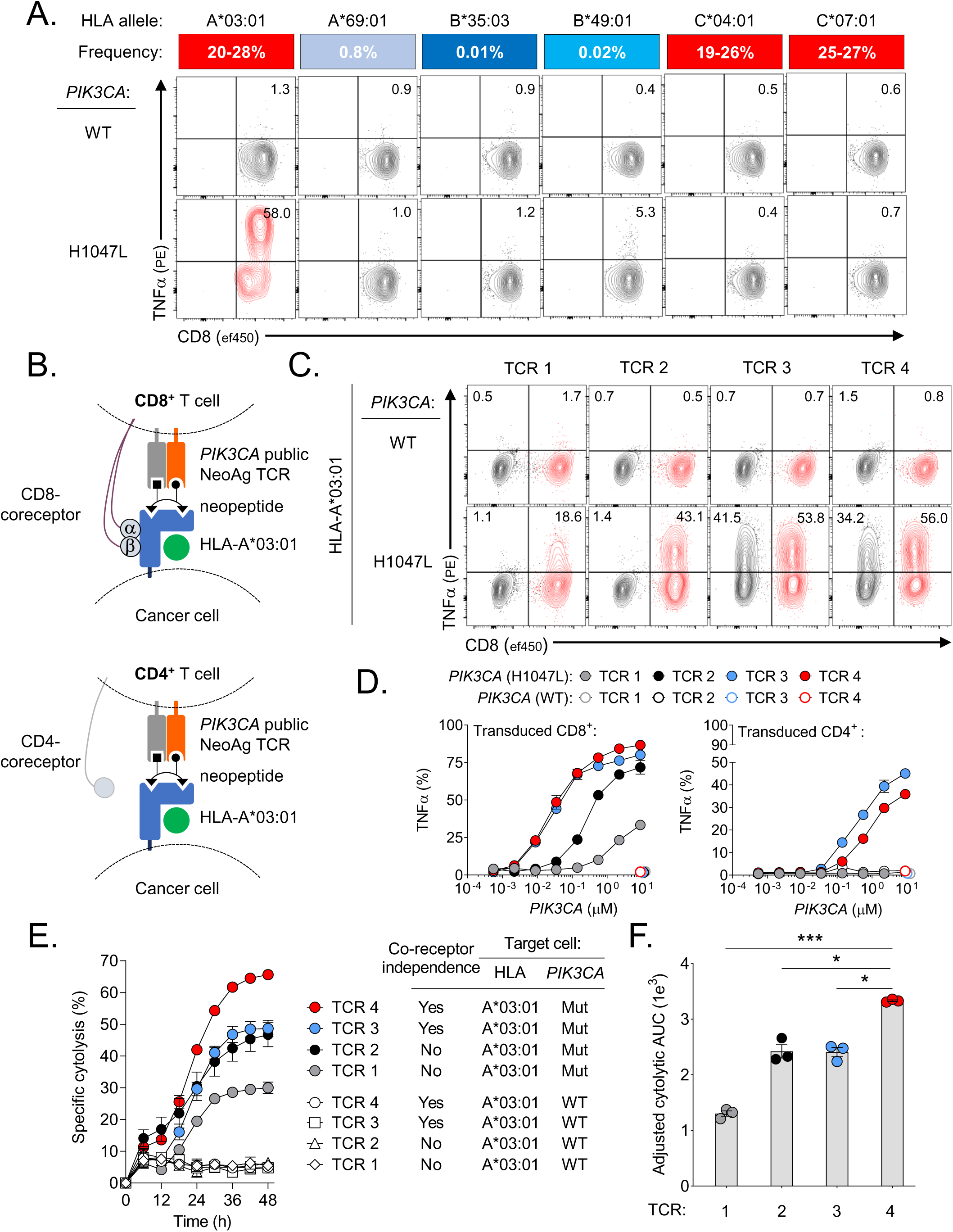
HLA restriction and functional characterization of *PIK3CA* public NeoAg-specific TCRs. (**A**) Deconvolution of the HLA class I (HLA-I) restriction element for the MSK 21LT2 clonotype 20 TCR. The frequency of individual HLA-I alleles expressed by HD1 in North American and European populations is displayed as a heat map. FACS plots show the frequency of CD8^+^ TCR transduced T cells that secrete TNFα following co-culture with HLA-I mono-allelic cell lines expressing WT or Mut *PIK3CA*. (**B**) Cartoon illustrating the experimental design to assess the co-receptor dependence and functionality of SIFT-seq retrieved *PIK3CA* public NeoAg TCR library members. TCRs 1-4 were individually retrovirally transduced into enriched CD8^+^ or CD4^+^ T cells and cocultured with target cells co-expressing A*03:01 and either WT or Mut *PIK3CA*. (**C**) Representative FACS plots of CD4^+^ (black) and CD8^+^ (red) T cells expressing individual PIK3CA public NeoAg TCR library members after co-culture with HLA-A*03:01^+^ target cells that express either WT or Mut *PIK3CA*. Numbers within each plot indicate the frequency of TNFα producing TCR transduced CD4^+^ (upper left quadrant) or CD8^+^ (upper right quadrant) T cells. The FACS plots shown in (**A**) and (**C**) are pre-gated on live^+^mTCR^+^ T cells. (**D**) The functional avidity of CD8^+^ (left) or CD4^+^ (right) T cells individually transduced with TCRs 1-4. Transduced T cells were co-cultured with an HLA-I mono-allelic cell line expressing HLA-A*03:01 and electroporated with indicated concentrations of WT or Mut *PIK3CA* mRNA. (**E**) Kinetic impedance-based lytic assay measuring the % specific cytolysis of A*03:01^+^ target cells expressing WT or Mut *PIK3CA*. (**F**) Adjusted cytolytic AUC values indicating the cumulative cytolytic capacity of TCRs 1-4 against A*03:01^+^/Mut *PIK3CA*^+^ target cells. All data shown is representative of ≥2 independent experiments using *n*=3 replicates per condition. Symbols and bar graphs displayed as mean ± SEM. * *P* ≤ 0.05; *** = *P* ≤ 0.001. Two-sided Student’s t-test with Bonferroni correction.

We next characterized the specificity, functional avidity, and cytolytic capacity of individual TCR library members to establish the relative potency of each receptor. In the context of HLA-A*03:01, all members exhibited absolute specificity for Mut *PIK3CA*; no responses were detected against WT *PIK3CA* (**Fig. 2C**). HLA-A*03:01 is the founding member of the HLA-A3 supertype, a group of homologous HLA-I molecules characterized by overlapping peptide-binding repertoires^26^. In select cases, different members of the HLA-A3 supertype can trigger cross-recognition by the same TCR^27,28^. Among the HLA-A3 supertype members, HLA-A*03:01 shares the highest sequence identity with HLA-A*03:02 (364/365 AA; 99.7%) and HLA-A*11:01 (358/365 AA, 98.1%). To determine if these related HLA-I alleles can trigger Mut-specific function, we co-cultured target cells expressing either HLA-A*03:01, -A*03:02 or -A*11:01 with T cells transduced with TCRs 1-4. All TCR library members solely recognized Mut *PIK3CA* in the context of HLA-A*03:01, highlighting each receptor’s exquisite selectivity for a single HLA-I restriction element (**Extended Data Fig. 2B** and **2C**).

We next compared the functionality of each TCR when transduced into isolated CD8^+^ and CD4^+^ T cells. The CD8 co-receptor binds to an invariant region present in all HLA-I molecules to stabilize sub-optimal interactions with the peptide/HLA-I complex typically observed with lower affinity TCRs^29^. When an HLA-I restricted TCR is transduced into a CD4^+^ T cell, the receptor must now function in the absence of this stabilizing interaction (**Fig. 2B**). The capacity of a TCR to function in a co-receptor independent manner has previously been correlated to a receptor’s intrinsic affinity to the peptide/HLA complex^29,30^ and clinical antitumor efficacy^31–34^. When expressed in CD8^+^ T cells, TCRs 3 and 4 triggered significantly greater Mut-specific cytokine production and degranulation compared with TCRs 1 and 2 (**Fig. 2C** and **Extended Data Fig. 3A**). When transduced into CD4^+^ T cells, we discovered that TCRs 3 and 4 were also capable of triggering T cell function, indicating co-receptor independence (**Fig. 2C**); by contrast, TCRs 1 and 2 were non-functional. Lastly, we quantified each TCR’s relative sensitivity to antigen abundance by determining their functional avidity. CD8^+^ and CD4^+^ T cells transduced with individual TCRs were co-cultured with HLA-A*03:01^+^ target cells expressing titrated quantities of either Mut or WT *PIK3CA* (**Fig. 2D**). In CD8^+^ T cells, TCRs 3 and 4 had lower EC_50_ values to Mut *PIK3CA* compared with TCRs 1 and 2 (**Extended Data Fig. 3B**); in CD4^+^ T cells, only TCRs 3 and 4 were functional.

Finally, we measured the capacity of CD8^+^ T cells transduced with TCRs 1-4 to mediate Mut-specific target cell killing using a kinetic impedance-based cytolytic assay. All four TCRs mediated Mut-specific cytolysis of a Mut *PIK3CA*^+^/HLA-A*03:01^+^ target cell but spared HLA-A*03:01^+^ cells expressing WT *PIK3CA* (**Fig. 2E**). Cytolysis was time-dependent for all TCRs, with maximum lysis measured at 48h. In repeated experiments, TCR4 exhibited superior lytic efficiency, as calculated by cytolytic area under the curve (AUC), relative to all other library members (**Fig. 2F**). We conclude that a public NeoAg derived from Mut *PIK3CA* can be targeted by TCRs with diverse functional attributes, including co-receptor independence and Mut-specific cytolytic function.

### Mechanism of immunogenicity for a PIK3CA public NeoAg

To resolve the mechanism of immunogenicity for the *PIK3CA* public NeoAg, we began by determining the sequences of endogenously processed and presented PI3Kα peptides presented by HLA-A*03:01. We developed a functional immuno-peptidomic screen using mono-allelic cell lines expressing individual HLA-A alleles and either Mut or WT *PIK3CA*. Peptide (p)/HLA-I complexes from these cell lines underwent immune precipitation/liquid chromatography tandem mass-spectrometry (LC-MS/MS). Skyline analysis revealed PI3Kα-derived amino acid (AA) sequences exclusively from cells that co-express HLA-A*03:01 and Mut *PIK3CA* (**Fig. 3A**). By contrast, no PI3Kα-derived peptides were observed from cells expressing HLA-A*03:01 and WT *PIK3CA* or Mut *PIK3CA* and an alternative HLA-A allele. Chromatogram analysis confirmed that all Mut PI3Kα-derived isotope precursor variants eluted at an identical retention time (**Fig. 3B**). Deconvolution of the MS/MS spectrum divulged a single 9 AA peptide containing the His→Leu substitution at the second position (P2) (A<u>L</u>HGGWTTK; henceforth pMut). The sequence of the eluted neopeptide was validated by matching its LC-MS/MS retention time and spectra with that of a synthetic peptide (**Fig. 3C**). Co-culture of peptide-pulsed HLA-A*03:01^+^ targets with Mut *PIK3CA*-specific T cells confirmed that only pMut but not its WT counterpart (A<u>H</u>HGGWTTK; henceforth pWT) triggered T cell function (**Fig. 3D and E**). Together, these data indicated that the LC-MS/MS-detected 9mer neopeptide encompassing the PI3Kα hotspot mutation was the minimal determinant responsible for T cell immunogenicity.

**Fig. 3:**
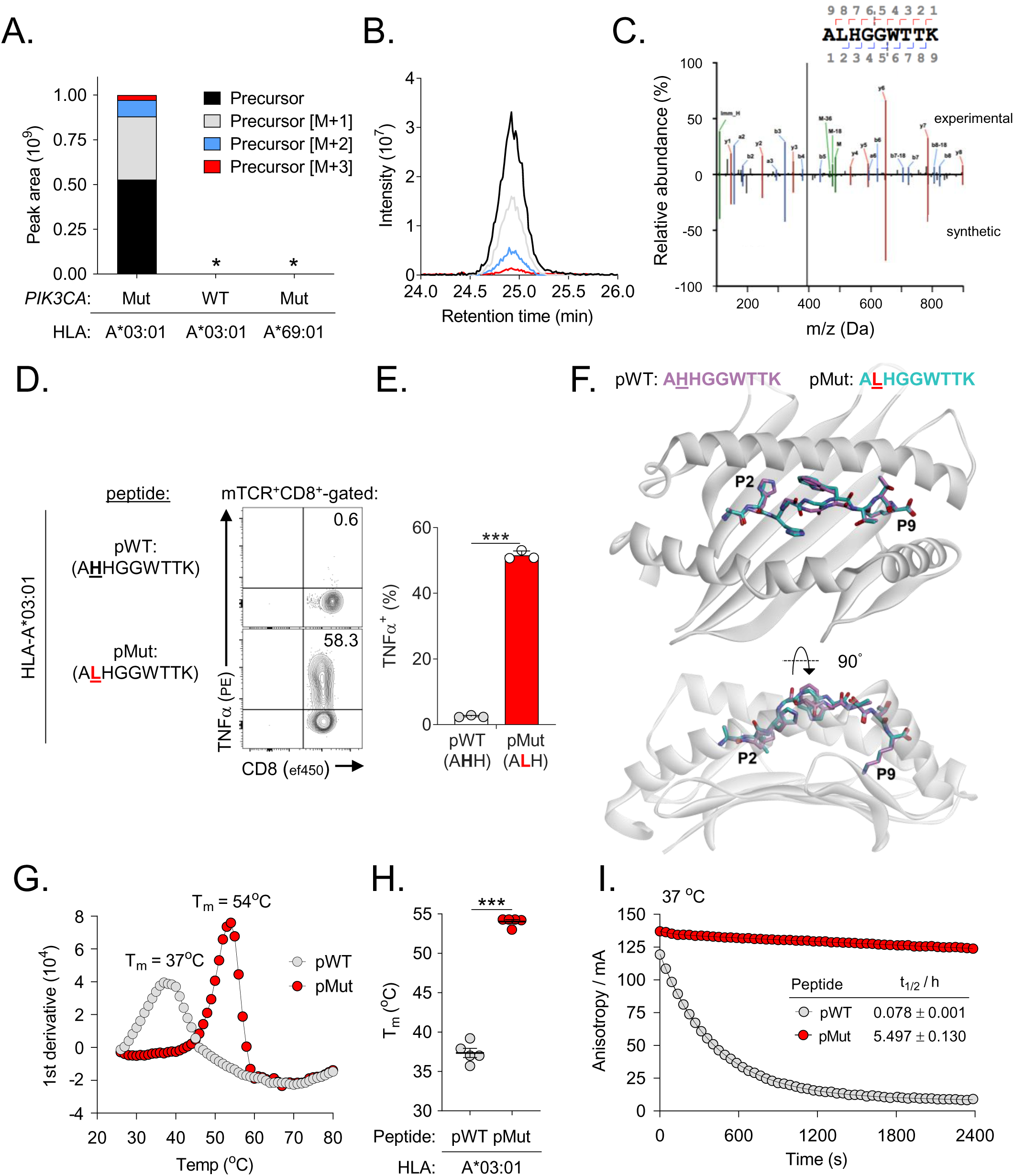
Mechanism of immunogenicity for a *PIK3CA* public NeoAg. (**A**) Skyline analysis of HLA-I bound peptides resulting from a LC-MS/MS-based immune-peptidomic screen. Relative abundance of precursor ions derived from PI3Kα protein eluted from HLA-A mono-allelic cell lines that endogenously express wild type (WT) or mutant (Mut) *PIK3CA* is shown. Asterix indicates conditions in which no PI3Kα-derived peptides were experimentally detected. (**B**) Corresponding chromatographic retention times of precursor ions derived from a Mut PI3Kα peptide eluted from HLA-A*03:01(A*03:01)^+^/Mut *PIK3CA*^+^ cells. (**C**) Mirror plot displaying the MS2 spectra of the Mut PI3Kα-derived public neoepitope (A<u>L</u>HGGWTTK; pMut; top) eluted from HLA-A*03:01^+^/Mut *PIK3CA*^+^ cells and a synthetically generated peptide (bottom). Peaks represent *b* ions in blue and *y* ions in red. (**D**) Representative intracellular FACS analysis and (**E**) summary bar graph for TNFα production in T cells transduced with TCR4 and co-cultured with HLA-A*03:01^+^ target cells pulsed with 1μM of pMut versus pWT. Results shown after gating on live^+^mTCR^+^CD8^+^ lymphocytes. Bar graph is displayed as mean ± SEM using *n*=3 replicates/condition. (**F**) Structural superimposition of the pMut and pWT peptides bound to HLA-A*03:01. The confirmations of pMut and pWT peptides are nearly identical with all α carbon atoms superimposing with a root mean square deviation (r.m.s.d.) of 0.73Å. (**G**) Representative thermal melt curves and (**H**) summary scatter plot displaying the melting temperatures (T_m_) of the pMut and pWT/HLA-A*03:01 complexes using differential scanning fluorimetry. Symbols are displayed as mean ± SEM using n=5 replicates. (**I**) Dissociation of a fluorescently-labeled pMut or pWT from soluble HLA-A*03:01 complexes at 37°C using fluorescence anisotropy. Solid lines show fits to exponential decay functions. Half-lives (t_1/2_) are shown for each peptide ± SEM. Data are representative of *n*=3. mA, millianisotropy. *** = *P* < 0.0001. Two-sided Student’s t-test.

We next sought to establish the physical basis for *PIK3CA* public NeoAg immunogenicity. After generating recombinant, soluble complexes, we crystallized and determined the structures of the pMut and pWT 9mer peptides bound to HLA-A*03:01 at resolutions of 2.0Å and 2.1Å, respectively (**Fig. 3F**; **Extended Data Fig. 4A-C**; **Supplementary Table 1**). Both peptides adopt conformations typical of nonameric peptides bound to HLA-I, bulging from the base of the HLA-A*03:01 peptide binding groove at positions 4-6. The geometries of the peptide binding grooves (residues 1-180 of the HLA-A*03:01 heavy chain) are nearly identical, with all α carbon (Cα) atoms superimposing with a root mean square deviation (r.m.s.d.) of 0.67Å. Notably, the conformations of the two HLA-I bound peptides are also nearly identical (Cα r.m.s.d. of 0.73Å). In both structures, the P6 Trp packs between the peptide backbone and the HLA-I α1 helix. Together with the P4 and P5 Gly residues, this results in a largely “featureless” central peptide region dominated by the peptide’s backbone. Similar to other cases where structures of WT and anchor-substituted p/HLA-I complexes have been compared^35^, the two peptides differ only slightly in their backbone conformations at the P4/P5 Gly residues (**Extended Data Fig. 4C**). This difference notwithstanding, the overall similarities of the pMut and pWT/HLA-I complexes suggested that structural differences alone cannot fully explain the immunogenic potential of the *PIK3CA* public NeoAg.

Recent studies indicate that the stability of a p/HLA-I complex has strong predictive value for private NeoAg immunogenicity^36,37^. Moreover, mutations that enhance peptide affinity through the creation of an optimal HLA-I anchor residue are also associated with immunogenicity^38–40^. We noted that the *PIK3CA* public NeoAg replaces a His with a Leu residue at P2; aliphatic AAs, especially Leu, are strongly preferred as a P2 primary anchor residue for HLA-A*03:01^41,42^. To quantify the relative stability of the pMut versus pWT peptides in complex with HLA-A*03:01, we performed differential scanning fluorimetry (DSF). DSF measures the melting temperature (T_m_) of a p/HLA-I complex, which serves as a proxy for peptide affinity^43^. Consistent with the introduction of an optimal P2 anchor, we discovered that the T_m_ of the pMut/HLA-I complex was significantly higher compared with the pWT/HLA-I complex (T_m_ 54°C Vs. 37°C; *P* < 0.0001; **Fig. 3G** and **3H**). This difference in T_m_ is similar in magnitude to other instances in which a P2-anchor modification leads to improved immunogenicity, including both shared tumor antigens and private NeoAgs^43,44^.

Improvements in peptide binding affinity are often associated with slower dissociation rates (measured as longer half-lives, t_1/2_). We therefore next resolved the kinetic stability of the two p/HLA-I complexes at physiologic body temperature (37°C) using fluorescence anisotropy (FA)^45^. The crystal structures of the pMut and pWT/HLA-A*03:01 complexes indicated that the P5 Gly side chains in both bound peptides point away from the HLA-I binding groove (**Extended Data Fig. 4C**). Consequently, we created versions of the pMut and pWT peptides fluorescently labeled at P5. We presumed this modification would be unlikely to influence the interaction of the peptide with the HLA-I protein (**Extended Data Fig. 4D**). At 37°C, the pMut/HLA-I complex had a significantly longer t_1/2_ compared to its pWT counterpart (t_1/2_ = 5.497h±0.130 Vs. 0.078h±0.001; **Fig. 3I**). Taken together, we conclude that the formation of the *PIK3CA* public NeoAg is driven by the creation of a peptide sequence containing an optimal HLA-I anchor residue. This results in a stable, high-affinity p/HLA-I complex with a prolonged t_1/2_ and demonstrable immunogenicity. By contrast, the pWT/HLA-I complex could not be detected on the surface of HLA-A*03:01^+^ cells, likely due to its instability at physiologic temperatures.

### Structural correlates of affinity and specificity for a PIK3CA public NeoAg TCR

A TCR’s specificity is primarily determined by the unique sequences of its six CDR loops, especially the somatically recombined CDR3 hypervariable loops^46^. We noted that TCR4 contains a CDR3β loop that is significantly longer (19 AAs) compared with TCRs 1-3 (range: 14-16 AAs) and a panel of >17,000 additional HLA-A*03:01 and -A*11:01-restricted TCRs (**Fig. 4A**)^47^. By contrast, the CDR3α AA length of TCR4 was within a comparable range to that of other TCRs. Because of TCR4’s unique CDR3β length and superior potency, we sought to characterize this receptor’s binding affinity, three-dimensional structure, and fine specificity.

**Fig. 4:**
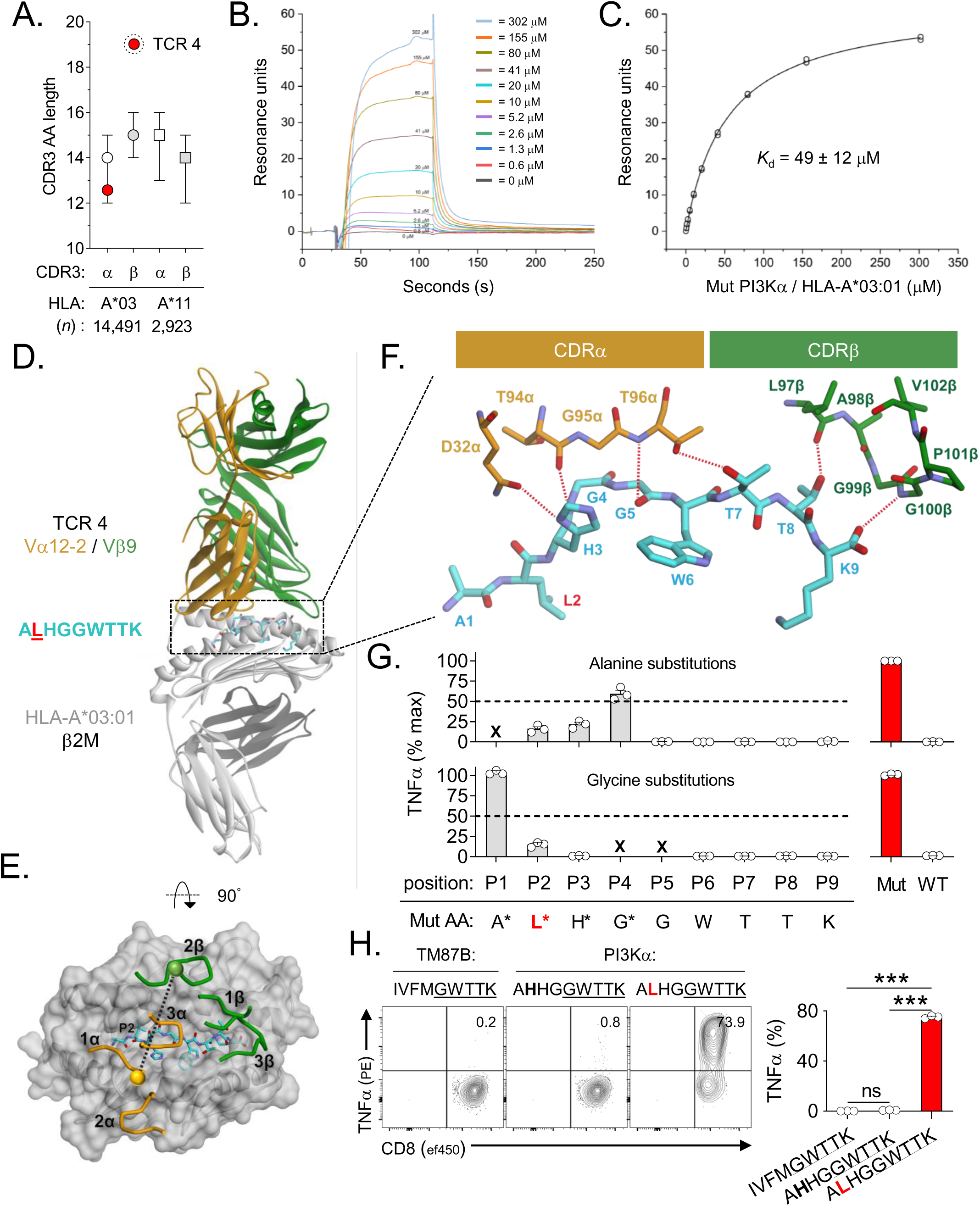
Structural correlates of affinity and specificity for a *PIK3CA* public NeoAg-specific TCR. (**A**) CDR3α and CRD3β amino acid (AA) lengths of *PIK3CA* public NeoAg-specific TCR4 and a panel of *n*=17,414 HLA-A*03:01 and -A*11:01-restricted TCR sequences. Results shown as median ± interquartile range. (**B**) Representative SPR sensorgram and (**C**) steady-state binding equilibrium measuring the dissociation constant (*K*_d_) for TCR4 to the pMut/HLA-I complex. Results shown are the average ± standard deviation of *n*=4 experiments. (**D**) Structural overview of the TCR4 pMut/HLA-A*03:01 ternary complex at 3.1Å resolution. The color scheme is indicated and replicated throughout. (**E**) Top view of the pMut/HLA-A*03:01 complex displaying the position of TCR4 and its crossing angle over the peptide/HLA-I along with the position of the six CDR loops. (**F**) Hydrogen bonds (red dashed lines) between TCR4’s CDRα and CDRβ chains and the pMut peptide. Amino acids are identified by standard one-letter codes followed by position number. The side chains of contacting residues from the TCR are also identified by AA and hemi-chain position number. (**G**) Identification of TCR4’s peptide recognition motif using alanine and glycine scanning. Intracellular FACS for TNFα production to Ala (upper) or Gly (lower) substituted peptides. TCR4-transduced T cells (identified by gating on mTCR^+^ lymphocytes) were cocultured with A*03:01^+^ targets pulsed with 1μM of indicated peptides. Results shown as the mean ± SEM percent maximum response relative to the native pMut peptide using *n*=3 biologic replicates per condition. “x” indicates positions not amenable to substitution. (**H**) Measurement of the cross-reactivity potential of TCR4. TCR4-transduced T cells were co-cultured with A*03:01^+^ targets pulsed with 1μM of peptides containing the motif “x-x-x-x-G-W-T-T-K”. Results shown as mean ± SEM using *n*=3 replicates per condition. *** = *P* < 0.0001. Two-sided Student’s t-test with Bonferroni correction.

We generated a recombinant, soluble version of TCR4 and measured its affinity towards the pMut/HLA-I complex using surface plasmon resonance (SPR), finding a *K*_d_ of 49(±12) μM (**Fig. 4B and C**). This affinity is higher than many TCRs specific for self-antigens and within a range typically observed for pathogen-specific receptors^48^. Using the same SPR experimental design at a temperature in which the pWT/HLA-I complex is stable (25°C; **Fig. 3G**), we failed to detect binding to the pWT/HLA-I complex (data not shown). This suggests that TCR4 might also distinguish between the subtle structural differences of pWT and pMut bound to HLA-A*03:01.

Pathogen-specific TCRs typically bind p/HLA-I complexes centrally and with a diagonal orientation^46^. This configuration positions the hypervariable CDR3α and CDR3β domains of the TCR over the peptide, contributing to both the affinity and specificity of a receptor. To explore how TCR4 binds the *PIK3CA* public NeoAg, we crystallized and determined the structure of the TCR4/pMut/HLA-A*03:01 ternary complex at a resolution of 3.1Å (**Fig. 4D**; **Supplemental Table 1**; **Extended Data Fig. 5**). The TCR docks using a conventional, diagonal orientation with an unremarkable crossing angle of 40° over the p/HLA-I complex (**Fig. 4E**). In most TCR structures, the two CDR3 loops pack against each other, centered around the pseudo-symmetry axis of the Vα/Vβ domains and focusing on the central regions of the peptide^49^. This is not the case with TCR4, however. The CDR3α loop is positioned parallel to the core of the “featureless” peptide backbone that extends from the P3 His to the P6 Trp (**Fig. 4F**; **Extended Data Fig. 5B,C**). The unusually long CDR3β loop, on the other hand, reaches out to the C-terminal end of the peptide, forming hydrogen bonds with the sidechain of the P8 Thr and the C-terminal carboxylate (**Fig. 4F**; **Extended Data Fig. 5A** and **B**). We further noted that TCR4 binding induces a significant change in the center of the peptide backbone, leading to a repositioning of the Trp6 side chain (**Extended Data Fig. 5C**). In the TCR bound state, Trp6 becomes fully buried in the HLA-A*03:01 peptide binding groove where it cannot form contact with the TCR. Thus, despite the pMut/HLA-I complex presenting a surface dominated by the peptide’s backbone, the unique structural properties of TCR4, and particularly its CDR3β loop, allow it to form a high complementarity interface that extends across nearly the entire peptide length.

Clinical application of high-affinity TCRs poses potential safety risks from on- and off-target reactivity^50,51^. We therefore assessed the specificity and cross-reactivity potential of TCR4 in greater detail. Using systematic AA replacements at each position of the cognate pMut peptide, we defined a consensus motif associated with this TCR’s function (**Fig. 4G**). T cells transduced with TCR4 were co-cultured with HLA-A*03:01^+^ target cells pulsed with Ala or Gly substituted peptides and the impact of these changes on cytokine production was determined. Replacement with either AA at positions P5-P9 resulted in near complete loss of function, indicating each residue’s critical role in TCR4 binding and recognition. Similarly, alterations in P2 and P3 of the peptide also led to a profound loss of cytokine secretion. Only substitutions at P1 and P4 were permitted. Collectively, these findings are consistent with the extensive interactions between TCR4’s CDR3α/β loops and the peptide, or in the case of the primary anchors at P2 and P9, the peptide and HLA-A*03:01. To retain a conservative threshold for establishing cross-reactivity potential, we selected all residues that permitted any measurable reactivity regardless of magnitude. Using this definition, we identified TCR4’s peptide recognition motif as “x-x-x-x-G-W-T-T-K” where “x” represents any AA. We next surveyed the human proteome for any sequence containing this motif using the ScanProsite tool^52^. This search afforded only two candidates: WT PI3Kα as expected and an unrelated peptide derived from transmembrane protein 87B (*TM87B*) with the sequence “I-V-F-M-G-W-T-T-K” (**Supplementary Table 2**). Importantly, the TM87B-derived peptide failed to elicit a cytokine response when pulsed onto HLA-A*03:01^+^ targets and co-cultured with TCR4 transduced T cells (**Fig. 4H**). Taken together, these data indicate that TCR4 binds the pMut/HLA-I complex similar to a pathogen-associated receptor and in a manner that maximizes CDR3 interactions with the neopeptide’s backbone. This configuration likely contributes to the receptor’s relatively high affinity and specificity for the *PIK3CA* public NeoAg.

### Clonality, immunogenicity, and immune resistance to a PIK3CA public NeoAg

NeoAg clonal heterogeneity has emerged as a major mechanism of resistance to T cell-based cancer immunotherapies^5–8^. Gain of function mutations in *PIK3CA* are critical to human cancer cell fitness^53,54^; we therefore hypothesized that *PIK3CA*(H1047L) would be an early evolutionary event and hence, clonally conserved across patients and disease sites. To test this hypothesis, we determined the cancer cell fraction (CCF), a measurement of mutation clonality, in a pan-cancer analysis of *n*=131 unique MSK patient samples harboring *PIK3CA*(H1047L). We discovered that Mut *PIK3CA* was clonally expressed in the majority of tumors (*n*=102/131; 77.9%), consistent with its reported role as a driver oncogene (**Fig. 5A**). Critically, clonal conservation of Mut *PIK3CA* was observed in both primary and metastatic tumor sites (**Fig. 5B**). Phylogenetic reconstruction using patient samples where matched primary and metastatic specimens were available revealed clonal preservation of Mut *PIK3CA* in the face of branched evolution for other mutated genes (**Fig. 5C**).

**Fig. 5:**
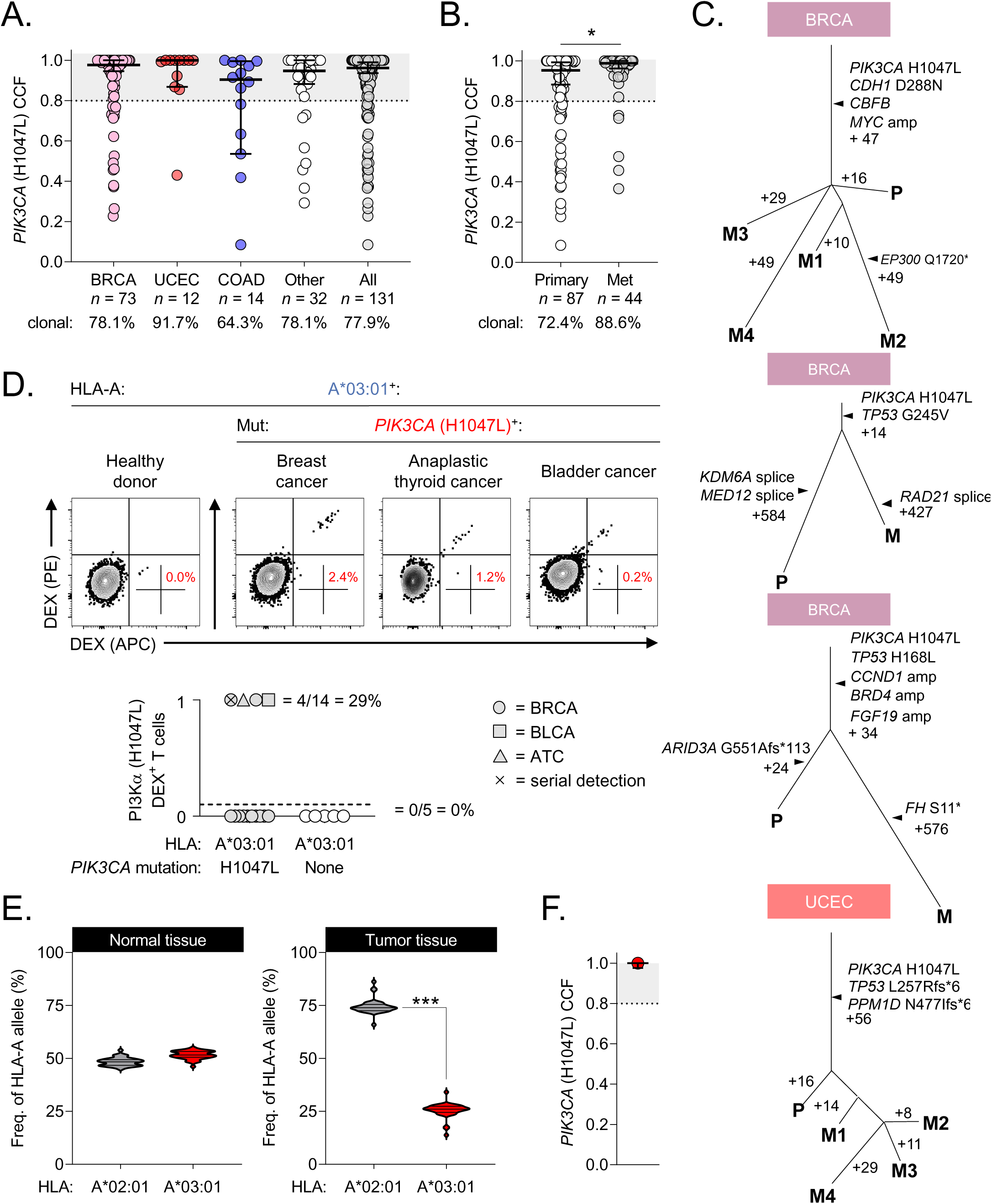
Clonality, immunogenicity, and immune resistance to a *PIK3CA* public NeoAg in cancer patients. **(A)** Pan-cancer analysis measuring the clonality of *PIK3CA* (H1047L) in *n*=131 unique MSK cancer patients. Clonality is defined as a cancer cell fraction (CCF) of ≥80%, indicated by the shaded grey area. (**B**) Comparison of *PIK3CA* (H1047L) CCF in primary versus metastatic tumor sites. (**C**) Phylogenetic analysis measuring the clonal conservation of *PIK3CA* (H1047L) in primary (P) and metastatic (M) tumor sites within the same patient. (**D**) Representative FACS and summary plot for the detection of circulating CD8^+^ T cells specific for the pMut/HLA-A*03:01 epitope in *n*=14 HLA-A*03:01^+^ cancer patients with a history of a *PIK3CA* (H1047L) cancer or *n*=5 HLA-A*03:01^+^ healthy donors. Percentages in FACS plots represent frequency of gated live^+^CD8^+^dual pMut/HLA-A*03:01 dextramer^+^ lymphocytes. 0 = no detection, 1 = detection. (**E, F**) Representative cancer patient who demonstrated a tumor-specific HLA-A*03:01 loss of heterozygosity in the setting of *PIK3CA* (H1047L) clonal conservation. Results in **A, B**, and **F** displayed as median ± 95% confidence interval. ATC = anaplastic thyroid cancer; BLCA = bladder cancer; BRCA = breast invasive carcinoma; COAD = colon adenocarcinoma UCEC = uterine corpus endometrial carcinoma. *** = *P* < 0.001; * = *P* < 0.05, two-sided Student’s t-test.

Having established the immunogenic potential and clonality of Mut *PIK3CA*, we hypothesized that a subset of HLA-A*03:01^+^ cancer patients would generate circulating immunity to this public NeoAg *in vivo*. To test this question, we developed a translational pipeline to prospectively collect PBMC samples from cancer patients of defined public NeoAg status (**Extended Data Fig. 6**). This effort was enabled by the MSK-Integrated Mutation Profiling of Actionable Cancer Targets (MSK-IMPACT) clinical next-generation sequencing (NGS) platform^55^. MSK-IMPACT performs NGS on both tumor and normal tissues, enabling the simultaneous detection of somatic tumor alterations and elucidation of germline encoded HLA-I alleles^56^. Using DARWIN, an automated genotype-driven enrollment tool that matches NGS results with upcoming patient clinic visits^57^, we flagged Mut *PIK3CA*^+^/HLA-A*03:01^+^ patients for biospecimen collection. This permitted us to curate a biorepository of PBMC samples from *PIK3CA* public NeoAg expressing cancer patients.

In total, we collected and analyzed samples from *n*=14 unique Mut *PIK3CA*^+^/HLA-A*03:01^+^ cancer patients (**Supplementary Table 3**). Cancer types in this cohort were diverse and representative of common malignancies associated with *PIK3CA* alterations. These include cancers of the breast (BRCA), endometrium (UCEC), colon (COAD), bladder (BLCA), high-grade gliomas (GBM), and anaplastic thyroid cancer (ATC)^19,58^. Using dual fluorochrome-labeled HLA-A*03:01 multimers loaded with the pMut neopeptide, we analyzed patient PBMC directly *ex vivo* and following a brief round of IVS. As controls, we simultaneously analyzed PBMC from HLA-A*03:01^+^ HDs using identical stimulation conditions. We detected circulating *PIK3CA* public NeoAg-specific T cells in 4/14 (29%) cancer patients and 0/5 HDs (0%) (**Fig. 5D**). In a subject with a history of Mut *PIK3CA* BRCA who is presently disease free and where serial blood samples were available, we consistently detected *PIK3CA* public NeoAg-specific T cells over time. This finding suggests that a patient can develop a sustained response to this public NeoAg. Together, these data support that the *PIK3CA* public NeoAg is immunogenic *in vivo* and is capable of driving T cell clonal expansion in a subset of Mut *PIK3CA*^+^/HLA-A*03:01^+^ cancer patients.

T cell targeting of immunogenic NeoAgs applies evolutionary pressure to cancer cells, potentially selecting for the outgrowth of populations with resistance to immune recognition^59^. Resistance can occur through multiple mechanisms^60^, including allele-specific HLA loss of heterozygosity (LOH)^61–63^. We therefore next sought to determine whether *PIK3CA* public NeoAg expressing tumors exhibited evidence of cancer cell-intrinsic immunologic resistance. Targeted exome sequencing did not reveal loss-of-function mutations in genes previously associated with resistance to T cell immunotherapies^64,65^, including *TAP1/2*, *CALR*, *B2M*, *HLA-A*, *JAK1/2*, and *IFNGR1*. By contrast, archival tissues from *n*=31 Mut *PIK3CA*^+^ /HLA-A*03:01^+^ cancer patients revealed HLA LOH involving the *A*03:01* allele in two subjects (2/31; 6.5%)(**Fig. 5E** and data not shown). No subjects exhibited LOH involving the non-A*03:01 allele. Interestingly, both subjects with HLA-A*03:01 LOH maintained clonal expression of *PIK3CA*(H1047L) (**Fig. 5F** and data not shown). Collectively, our findings demonstrate that the *PIK3CA* public NeoAg is typically clonally conserved, immunogenic in cancer patients, and in select cases associated with a mechanism of immune resistance.

## DISCUSSION

Here, we demonstrate that a hotspot mutation in PI3Kα, among the most commonly altered driver genes^19,20^, is endogenously processed, presented, and immunogenic in the context of a prevalent HLA-I allele, thereby creating a public NeoAg. Unlike private NeoAgs resulting from patient-specific passenger mutations, we discovered that the *PIK3CA* public NeoAg is both shared among patients and frequently clonally conserved. Using SIFT-seq, a functional TCR discovery platform that is agnostic of HLA binding predictions, we generated a library of unique TCR gene sequences that confer specificity to this shared NeoAg. These data establish the immunogenic potential of the *PIK3CA* public NeoAg, enabling future off-the-shelf therapies using TCR engineered T cells or antigen-specific vaccination.

Mechanistically, we determined that the molecular basis for *PIK3CA* public NeoAg immunogenicity results primarily from the creation of an optimal HLA-I anchor residue formed by the H1047L hotspot mutation. Crystal structures of the pMut and pWT/HLA-I complexes revealed that the peptides adopt very similar conformations in the HLA-A*03:01 binding groove, both presenting a largely “featureless” peptide backbone. The similarity of the pMut and pWT/HLA-I complexes thus suggest that PIK3CA NeoAg immunogenicity arises from a separate mechanism. Aliphatic AAs, especially Leu, are strongly preferred in the P2 anchor position of A*03:01^26,42^. We therefore hypothesized that pMut would preferentially stabilize the HLA-I complex compared with pWT. Consistent with our hypothesis, we found that the kinetic stability of the pMut/HLA-I complex was substantial at physiologic temperature, which *in vivo* would result in sustained antigen presentation. In contrast, the pWT/HLA-I complex rapidly degraded, a finding that corresponds with our inability to detect pWT on the surface of HLA-A*03:01^+^ cells by mass spectrometry. These findings are consistent with several recent reports that identified neopeptide/HLA-I complex stability as a major determinant for private NeoAg immunogenicity in humans^36,37^.

Prior studies indicate that co-receptor independence and lytic capacity correlate with clinical efficacy for TCR gene therapies targeting self^31^, over-expressed^66^, viral^34^, and cancer-germline antigens^33^. Stratification of our SIFT-seq retrieved TCR library members revealed that TCR4 was co-receptor independent, functionally avid, and capable of sustained cytolysis of Mut *PIK3CA*^+^/HLA-A*03:01^+^ target cells. TCR4 possesses a relatively high affinity (*K*_d_ ∼50μM) that is within a range associated with pathogen-specific^48^ and co-receptor independent TCRs^29^. Despite the receptor’s affinity, we did not detect binding of TCR4 to the pWT/HLA-I complex, even at reduced temperatures. This suggests that the receptor can distinguish subtle structural differences of pWT and pMut bound to HLA-A*03:01, mirroring previous findings using altered peptide ligands for TCRs specific for self and viral antigens^67,68^. Our data, together with recently published structural studies from a limited number of additional NeoAg-specific TCRs^69–71^, indicate that the human T cell repertoire can perceive mutated self-proteins in a manner analogous to pathogen-associated determinants. By contrast, many self-antigen specific TCRs are co-receptor dependent and possess comparatively weaker affinities (>50μM), likely reflecting the influence of negative thymic selection^48^.

Comparison of TCR4’s gene sequence to other library members and a panel of >17,000 additional TCRs restricted by A*03:01 or A*11:01 revealed this receptor possesses an unusually long CDR3β length. The functional significance of TCR4’s long CDR3β loop was revealed by resolving the structure of the receptor in complex with pMut/HLA-I. The CDR3 loops of most TCRs engage the central core (P4-P7) of an HLA-I bound peptide. However, the PIK3CA pMut/HLA-I complex presents a challenge: the surface it displays to a TCR is relatively “bland” because it is dominated by the peptide’s backbone rather than exposed side chains typically associated with peptide immunogenicity^72^. Indeed, upon TCR4 binding, the side chain of the neopeptide’s most distinctive residue (P6 Trp) becomes fully buried in the HLA-A*03:01 peptide binding groove, further increasing its featureless nature. Recent efforts to rationalize the importance of bulky, multifunctional amino acids such as Trp in peptide immunogenicity have emphasized their importance in forming key TCR contacts^72^. This principle was recently illustrated in the TCR recognition of an ovarian cancer-associated private NeoAg^69^. TCR4 recognition of the *PIK3CA* public NeoAg clearly defies such expectations.

To solve the challenge, TCR4’s unusually long CDR3β loop enables the receptor to form an extended and highly complementary interaction with the peptide, including important interactions with its two most C-terminal residues. This unusual and distinctive configuration likely contributes not only to TCR4’s affinity but also its specificity as substitution of native amino acids in 7-9 peptide positions results in near complete loss of function. Consequently, TCR4 was found to lack measurable cross-reactivity to any alternative epitope derived from the human proteome. The exquisite fine specificity of TCR4, combined with: 1) the tumor-specific expression of Mut *PIK3CA*, and 2) the absence of a stable WT p/HLA complex on normal tissues, establishes a favorable safety profile for this receptor’s future clinical development. Understanding how TCRs with conventional CDR3 lengths engage the pMut/HLA-I complex, including whether they differentially engage invariant regions of HLA-A*03:01, will be an important area of future research.

Circulating T cells specific for the *PIK3CA* public NeoAg were detected in ∼30% of HLA-A*03:01^+^ patients with a history of a Mut *PIK3CA*^+^ cancer but 0% of matched HLA-A*03:01^+^ HDs. This finding suggests that endogenous levels of Mut PI3Kα is immunogenic *in vivo* and capable of driving the clonal expansion of public NeoAg-specific T cells. T cell-mediated attack on cancer cells can apply evolutionary selection pressure resulting in the outgrowth of tumor cells harboring genomic alterations that confer immune-resistance^59,60^. Loss of heterozygosity for the restricting HLA-I allele of an immunogenic NeoAg is a precise means of immune-editing that does not compromise tumor cell fitness^61–63^. We found selective LOH for HLA-A*03:01, but not alternative HLA-A alleles, in 2/31 (7%) of patients. Expression of Mut *PIK3CA* was clonally conserved in both of these subjects. Similar findings were recently reported using an independent pan-cancer data set of >83,000 HLA-I typed patients: in those subjects who were HLA-A*03:01^+^ and harbored a *PIK3CA*(H1047L)^+^ cancer, there was a statistically significant LOH for HLA-A*03:01^+^ over the alternative HLA-A allele^63^. Together, these data suggest that the immunogenicity of *PIK3CA* public NeoAg expressing tumors may lead to acquired immune-escape while preserving oncogenic driver expression. Across our entire MSK cohort of public NeoAg expressing cancers, targeted exome sequencing did not reveal alternative genomic alterations that might compromise antigen processing and presentation. However, this finding does not preclude tumor cell-intrinsic transcriptional^73^ and/or epigenetic^59^ mechanisms of antigen presentation immune escape which have also recently been reported. We have assembled what is to our knowledge the largest collection of biospecimens from patients who express an identical NeoAg resulting from a NSSM. Nevertheless, a limitation of our study is that the sample size remains underpowered to conclusively establish correlates between patient demographics, tumor-specific parameters, and immunogenicity.

In conclusion, we report on the development of an innovative new platform to identify and retrieve TCRs specific for shared NeoAgs resulting from mutated driver genes. Using this platform, we identified an immunogenic public NeoAg derived from Mut *PIK3CA*, resolved its mechanism of immunogenicity, and established a library of cognate TCRs that could be deployed as an “off-the-shelf” therapy, with one TCR in particular demonstrating considerable promise. However, the modular nature of the SIFT-seq platform might readily be applied to the discovery of public NeoAg-specific TCRs resulting from the >200 additional driver genes identified to date^19^. Finally, we demonstrate a novel application of clinical NGS to efficiently identify cancer patients who express a requisite HLA-I allele and Mut driver oncogene to create a public NeoAg. Because TCR-based therapeutics require knowledge of two biomarkers (target antigen expression and HLA allele), this approach overcomes a key bottleneck currently limiting clinical dissemination. Recent regulatory approvals have been made for tissue-agnostic therapies targeting specific genomic alterations identified using clinical NGS, including NTRK fusion inhibitors^74,75^ and immune checkpoint inhibitors in microsatellite unstable cancers^76^. An analogous approach might be applied for the future clinical development of target-specific but tissue-agnostic TCR therapies targeting public NeoAgs.

## METHODS

### Primary cells and cell lines

Leukopaks from HD were purchased from the New York Blood Center (NYBC). HLA-typed leukoreduction system (LRS) chambers from HDs were obtained from the Stanford Blood Center. PBMCs were isolated by density-gradient centrifugation using lymphocyte separation medium (Corning, REF 25-072-CV) and cryopreserved until ready for use. High resolution genomic *HLA*-typing for HD PBMC was performed by Histogenetics. Patient-derived PBMC were obtained after signed informed consent from CPT collections drawn during scheduled clinical appointments. Protocols were reviewed and approved by the Institutional Review Board (IRB) at MSKCC (IRB protocols 17-245 and 17-250). The retroviral packaging line 293-GP was purchased from Takara Bio. COS-7 cells were obtained through an MTA from S. A. Rosenberg (NCI, Bethesda, USA). Human primary cells were cultured in complete RPMI 1640 (Gibco, REF 11875-085) media supplemented with antibiotics and 10% human AB serum (Gemini Bioproducts, Cat: 100-512). Cell lines were cultured and maintained in RPMI 1640 (Gibco) media supplemented with Pen/Strep (Gibco, REF 15070-063), Gentamycin (MP Biologicals, Cat 1676245) and 10% FBS (Gemini Bioproducts, Cat 100-106).

### Plasmids and peptides

All *PIK3CA* and *HLA* sequences were synthesized and cloned into the pcDNA3.1^+^ vector (Genscript). Constructs encoding WT *PIK3CA* and individual *PIK3CA* hot spot mutations (E542K, E545K and H1047L/R) were generated. Plasmid vectors were linearized using XbaI (New England Biolabs, Cat R0145S) and purified using QIAquick PCR purification kit (Qiagen, Cat 28104) and following manufacturer’s instructions. *In vitro* transcribed RNA was generated using the HiScribe T7 High Yield RNA Synthesis kit (New England Biolabs, Cat E2060S) following manufacturer’s instructions. TCR α and β sequences were synthesized and cloned into the pMSGV-1 retroviral plasmid vector as previously described^77^. The human variable (V) regions of the retrieved TCRs were fused to a murine constant chain (mTCR) to facilitate identification of transduced T cells and promote proper chain pairing^78^. HPLC grade 9-mer peptides (>99%) for pWT, pMut, and a peptide derived from TM87B were manufactured by Genscript.

### *In vitro* sensitization (IVS) of HD and patient PBMC

PBMCs were plated in tissue culture flasks at 1e^6^ cells/cm^2^ in complete media in the absence of cytokine for 2h at 37°C to separate the adherent (monocyte-containing) and non-adherent (T cell containing) fractions. CD8^+^ and CD4^+^ cells were enriched from the non-adherent fraction by untouched negative selection (Stemcell Technologies, Cat 17952 and Cat 17953). To generate moDC, the adherent fraction was washed with PBS and fresh complete media supplemented with recombinant human IL-4 and GM-CSF (400 IU mL^−1^) was provided every alternate day. moDC were stimulated with LPS (Invitrogen, REF 00-4976-93) and IFN-γ (Miltenyi Biotec, Cat 130-096-484) for 16-24h prior to transfection. moDCs are electroporated with 100 μg mL^−1^ mRNA encoding an individual PIK3CA hotspot mutation. HD T cells were co-cultured with electroporated moDCs at an E:T ratio of 3:1 in the presence of IL-21 (30 ng mL^−1^) in 24 well non-TC plates (Falcon, 351147). Wells were supplemented with fresh media containing IL-7 and IL-15 (10 ng mL^−1^ each) every 3 days during the duration of the *in vitro* culture. All cytokines were purchased from Miltenyi Biotec. For patient-derived samples, PBMCs were enriched for CD8^+^ T cells and stimulated with autologous monocytes pulsed with p/Mut (1μg mL^−1^) in the presence of 300 IU mL^−1^ IL-2.

### Mutation-specific qPCR screen

Aliquots containing 2e^4^ cells from individual parent IVS wells were harvested, split equally into two daughter wells, and stimulated with moDCs electroporated with mRNA encoding WT or Mut *PIK3CA* for 3h at an E:T ratio of 1:2. Post-stimulation, total mRNA was purified (Qiagen), transcribed into cDNA (ABI, Ref 4368813) and levels of *IFNG* transcript are quantified by real-time qPCR (ABI QuantStudio 5) using the TaqMan Fast Advanced MasterMix (ABI, Cat 4444557). Delta CT values for matched Mut versus WT stimulated wells were calculated. Wells with a positive delta CT value ≥ 2SD from the mean were selected for downstream single-cell immune-profiling.

### Single-cell immune-profiling

Single-cell RNA and V(D)J sequencing was performed using the 10x Genomics platform. 1e10^5^ cells from a qPCR-positive well were harvested and co-cultured with moDCs electroporated with mRNA encoding WT or Mut *PIK3CA* at an E:T ratio of 1:10 for 3h. Final cell concentration was determined by cell counting on a hemocytometer, and cell concentration was adjusted to approximately 700 and 1,200 cells μL^−1^ to maximize the likelihood of achieving the desired cell recovery target, with an initial cell viability of >90%. The single-cell suspension was mixed with RT Master Mix and loaded together with barcoded single-cell 5′ gel beads and partitioning oil onto Single Cell A Chip to generate GEMs using Chromium Controller. Cell lysis and barcoded reverse transcription of RNAs from single cells were finished inside each GEM. Barcoded complementary DNA (cDNA) product was recovered through post GEM-RT cleanup and PCR amplification. cDNA QC and quantification were determined by High Sensitivity D5000 DNA ScreenTape analysis (Agilent Technologies) and Qubit dsDNA HS Assay Kit (Thermo Fisher Scientific). Fifty nanograms of cDNA was used for 5′ Gene Expression library construction, and each sample was indexed by a Chromium i7 Sample Index Kit, which was run on an Illumina HiSeq4000 sequencer with 2 × 100 base pairs (bp) paired reads to achieve at least 30,000 read pairs per cell. Full-length V, D and J gene segments were amplified using a Chromium Single Cell V(D)J Enrichment Kit (human T cells) to generate enrichment products. Enriched product was measured by D5000 DNA ScreenTape analysis and Qubit dsDNA HS Assay Kit, and then 50 ng of enrichment TCR product was used for library construction. Single Cell V(D)J enriched libraries were sequenced on HiSeq4000 to produce paired 2 × 150 bp reads at 5,000 read pairs per cell. The raw scRNA-seq and VDJ data were pre-processed (demultiplexing of cellular barcodes, read alignment and generation of feature-barcode matrix) using Cell Ranger (10x Genomics, v2.1.1) with the annotation files “vdj_GRCh38_alts_ensembl-3.1.0-3.1.0” and “GRCh38-3.0.0” for the V(D)J sequencing and gene expression libraries, respectively. Detailed QC metrics were generated and evaluated using single-cell analysis packages from Bioconductor including DropletUtils, scater, and scran. Genes detected in fewer than three cells and cells where fewer than 200 genes had nonzero counts were filtered out and excluded from subsequent analysis. Low-quality cells where more than 15% of the read counts derived from the mitochondrial genome were also discarded. For downstream analyses, only cells for which clonotype information was available were retained. To remove likely doublet or multiplet captures, cells with more than 7,000 detected genes were discarded as well as cells for which more than 2 *TRA* or *TRB* sequences were detected. For SIFT-seq based identification of (i) T cell clonotypes associated with a Mut *PIK3CA*-specific response, (ii) gene expression pattern analyses of candidate clonotypes, and (iii) visualizations and TCR retrieval, custom R scripts were developed.

### HLA-IP LC-MS/MS

COS-7 cells were co-electroporated with 100 μg mL^−1^ each of mRNA encoding an individual *HLA* allele with either WT or Mut *PIK3CA*. 15-20e10^6^ cells were electroporated per condition and plated in 6 well plates overnight. Cells were harvested by incubating with 1mM EDTA (Millipore Sigma, Cat 324504). Harvested cells were pelleted and washed 3 times in ice-cold sterile PBS (Gibco, Cat 10010-0230. Immunoprecipitation, HLA ligand separation and LC-MS/MS were performed as previously described^79^. Briefly, cells were lysed in 7.5 mL of 1% CHAPS (MilliporeSigma) for 1h at 4°C, lysates were spun down for 1h with 20,000*g* at 4°C, and supernatant fluids were isolated. For immune-purification of HLA-I ligands, 0.5 mg of W6/32 antibody (BioXCell) were bound to 40 mg CN-Br activated sepharose and incubated with the protein lysate overnight. HLA complexes and binding peptides were eluted five times using 1% TFA. Peptides and HLA-I complexes were separated using C18 columns (Sep-Pak C18 1 cc Vac Cartridge, 50 mg sorbent per cartridge, 37–55 μm particle size, Waters). C18 columns were preconditioned with 80% ACN (MilliporeSigma) in 0.1% TFA and equilibrated with 2 washes of 0.1% TFA. Samples were loaded, washed again twice with 0.1% TFA, and eluted in 300 μL of 30%, 40%, and 50% acetonitrile in 0.1% TFA. All three fractions were pooled, dried down using vacuum centrifugation and stored at −80°C until further processing. HLA-I ligands were isolated by solid-phase extractions using in-house C18 minicolumns (REF). Samples were analyzed by high-resolution/high-accuracy liquid chromatography-tandem mass spectrometry (LC-MS/MS) (Lumos Fusion, ThermoFisher Scientific). MS and MS/MS were operated at resolutions of 60,000 and 30,000, respectively. Only charge states 1, 2, and 3 were allowed. The isolation window was chosen as 1.6 thomson, and collision energy was set at 30%. For MS/MS, maximum injection time was 100 ms with an automatic gain control of 50,000. MS data were processed using Byonic software (version 2.7.84, Protein Metrics) through a custom-built computer server equipped with 4 Intel Xeon E5-4620 8-core CPUs operating at 2.2 GHz and 512 GB physical memory (Exxact Corporation). Protein FDR was disabled to allow complete assessment of potential peptide identifications. Oxidization of methionine; phosphorylation of serine, threonine, and tyrosine; as well as N-terminal acetylation were set as variable modifications for all samples. Samples were searched against a database comprising UniProt *Cercopithecus* Aethiops reviewed proteins supplemented with human HLA allele sequences used in this study, WT and Mut sequences of PI3Kα as well as common contaminants. For mirror plots ion intensities were exported and re-plotted with Graphpad prism. For precursor quantitation as well as retention time analyses skyline software (version 4.2, MacCoss Lab Software) was used. Precursor masses of peptide target sequences were searched in all relevant .raw files and peak areas of all replicates compared. Retention times for the best scoring matches were selected with a total of 4 isotopes.

### TCR transduction

293GP packaging line was plated O/N into poly-D-lysine coated 60mm^2^ plates at 1.6e^6^ cells/ plate. 293GP cells were then transfected with 6 μg of a candidate TCR pMSGV1-plasmid in the presence of the 3 μg of RD114 envelope using Lipofectamine 3000 (Invitrogen). The pMSGV1-plasmid and RD114 were obtained through an MTA from S. A. Rosenberg (NCI, Bethesda, USA). Viral supernatants were collected after 48h and loaded onto retronectin-coated (10 ug mL^−1^, Takara Bio) non-TC 24-well plates. Open-repertoire T cells (7.5e10^6^ cells per well of a 24 well plate) were stimulated with OKT3 (50 ng mL^−1^, Miltenyi Biotec, Cat 130-093-387) and rhIL-2 (300 IU mL^−1^) for 48-72h prior to transduction, and transduction was carried out as described previously (Hughes 2005, Human Gene Therapy). At 3-4d post-transduction, cells were assessed for tranduction efficiency by measuring surface expression of mTCR by FACS.

### T cell immunoassays

<u>Mutation-specificity</u>: Specificity of candidate TCRs was assessed by intracellular cytokine staining (ICS) using the BD Cytofix/CytoPerm Plus kit (Cat 555028) following manufacturer’s instructions. moDCs from MSK21LT2 and 0606T donors were electroporated with 100 μg mL^−1^ IVT mRNA encoding WT or Mut *PIK3CA* and plated into 96-well round bottom plates O/N. To determine the class of MHC restriction, a set of plated moDCS were treated with an anti-HLA-A/B/C antibody (40 μg mL^−1^, Clone W6/32, Biolegend Cat 311428) or anti-class II (10 μg mL^−1^, Clone IVA12, MSKCC in-house) antibodies for 3h at 37°C, as previously described. Candidate TCR-transduced T cells were then added in at an E:T ratio of 1:1 for 6h in the presence of anti-CD107A-BV650 (Clone H4A3, Biolegend 328638) and Golgi Block. PMA-ionomycin stimulation is included as a positive control. Following 6h of coculture, cells were washed in 1x PBS and surface labeled with Live/Dead fixable dye (Invitrogen), anti-CD3-APC-H7 (Clone SK7, BD 641397), anti-CD4-Alexa Fluor 700 (Clone RPA-T4, Invitrogen 56-0049-42), anti-CD8-efluor450 (Clone SK1, Invitrogen 48-0087-42) and anti-mouse TCR-PerCpCy5.5 (Clone H57-597, Invitrogen 45-5961-82) for 30 min at 4°C. Cells were washed with 1x PBS and then fixed and permeabilized for 15’ at 4°C. Surface-labeled cells were then washed with 1x perm-wash buffer and labeled with anti-IL-2-PE-Cy7 (Clone MQ1-17H12, Invitrogen 25-7029-42), anti-TNFα-PE (Clone Mab11, Invitrogen 12-7349-82) and anti-IFN-γ-FITC (BD 554551) for 30 minutes at 4°C in perm-wash buffer. Finally, cells were washed with perm-wash buffer and suspended in 2% FBS in PBS prior to acquisition on the X20 LSR Fortessa flow cytometer. Data were analyzed using FlowJo software version 10.6.2. A representative example of a FACS gating strategy is shown in **Extended Data Fig.7**. <u>HLA-restriction determination:</u> COS-7 cells were co-electroporated with 100 μg mL^−1^ per mRNA encoding an individual *HLA* allele with either WT or Mut *PIK3CA*. Transfected COS-7 cells are plated overnight in a 96-well round bottom plate in complete media to allow surface p/HLA-I complex expression. TCR-transduced T cells are added in at an E:T ratio of 1:1 and cytokine production was measured by ICS as described above. <u>TCR coreceptor dependence and functional avidity</u>: Target cells expressing HLA-A*03:01 were electroporated with mRNA encoding Mut or WT *PIK3CA* and plated overnight. Enriched CD4^+^ or CD8^+^ expressing an individual Mut *PIK3CA*-specific TCR are added in at a 1:1 E:T ratio. EC_50_ values for individual TCR are calculated by determining antigen concentration that elicits 50% of maximal response. <u>TCR killing assay</u>: TCR cytolytic capability was measured by a tumor impedance assay using the xCELLigence system (ACEA Biosciences). Target cells expressing HLA-A*03:01 and either Mut or WT *PIK3CA* were plated overnight into custom xCELLigence E-plate 96-well flat bottom plates. Prior to adding T cells, a baseline impedance measurement was taken. TCR-transduced T cells were added at an E:T ratio of 1:4 and impedance measurements were recorded at 15 min intervals for up to 96 h. Targets plated in complete media or exposed to 1% Triton-X allowed minimum and maximum lysis measurement, respectively. Percent cytolysis was calculated using the RTCA Software Pro. <u>TCR4 cross-reactivity</u>: Target cells expressing HLA-A*03:01 were pulsed with WT or Mut PI3Kα 9-mers or variants that contain an alanine or glycine substitution at each individual peptide position (1 μg mL^−1^) for 60 min at 37°C. Unbound peptide was washed off and TCR4-transduced T cells were added at an E:T ratio of 1:1. Residues required for TCR4 function are identified to derive a peptide recognition motif. ScanProsite was used to search all UniProtKB/Swiss-Prot database sequences, including splice variants, for proteins containing the motif “X-X-X-X-G-W-T-T-K”. No filters were used.

### Dextramer labeling

HLA-A*03:01 multimers bound to Mut PI3Kα peptide and conjugated to PE or APC were purchased from Immudex. Cells were labeled with dual fluorophore-conjugated dextramers for 10 min at RT followed by surface antibodies against CD3, CD4, CD8 and viability dye for an additional 20 min at 4°C. Cells are washed and acquired on the BD Fortessa X20 flow cytometer.

### Recombinant protein synthesis and isolation

Proteins for biophysical assays and crystallography, including the HLA-A*03:01 heavy chain, β_2_-microglobulin (β_2_m) and the TCR α/β chains were expressed in *Escherichia coli* as inclusion bodies and dissolved in 8M urea and 6M guanidinium-HCl. Denatured proteins were refolded and purified *in vitro* as previously described^80^. Briefly, for TCR folding, TCR α/β chains at a 1:1 ratio were diluted into TCR refolding buffer (50 mM Tris-HCl, 2.5 M urea, 2 mM NaEDTA, 6.5 mM cysteamine, 3.7 mM cystamine and 0.2 mM PMSF, pH 8.15) and incubated at 4°C overnight. The refolding buffer was then dialyzed against ddH_2_O and 10 mM Tris-HCl (pH 8.3) at 4°C for 36 h. For peptide/MHC folding, human β_2_m and HLA-A*03:01 were injected into the refolding buffer (400 mM L-arginine, 100 mM Tris-HCl, 2 mM NaEDTA, 6.3mM cysteamine, 3.7mM cystamine and 0.2 mM PMSF, pH 8.30), respectively, at a 3:1 ratio in the presence of a 10-fold excess of peptides at 4°C. After an overnight incubation, the solution was dialyzed against ddH_2_O and 10 mM Tris-HCl (pH 8.3) at room temperature for 48 h. The refolded proteins were then purified via ion-exchange followed by size-exclusion chromatography. Protein concentrations were determined through UV absorbance using sequence-determined extinction coefficients. WT and Mut PI3Kα peptides were purchased from AAPPTEC or GenScript at >80% purity, diluted to 30 mM in DMSO, and stored at −80°C.

### Differential scanning fluorimetry

Melting temperatures of WT and Mut PIK3α peptide/HLA-A*03:01 complexes were measured via differential scanning fluorimetry (DSF) on a StepOnePlus Real-Time PCR System (Applied Biosystems) as previously described^43^. In brief, a 96-well plate was loaded with mixtures of 2 μL 100X SYPRO Orange protein gel stain (Invitrogen) and 18 μL target protein at 15 μM in HBS-EP buffer (10 mM HEPES, 150 mM NaCl, 3 mM NaEDTA, 0.005% surfactant P20, pH 7.4). The excitation and emission wavelengths for the measurement were 587 nm and 607 nm, respectively. Temperature was scanned from 20°C to 95°C at a rate of 1°C min^−1^. Melting temperatures were determined by fitting the 1^st^ derivative of each melting curve to a bi-Gaussian function in OriginPro.

### Fluorescence anisotropy

Dissociation of WT and Mut PI3Kα peptides from peptide/HLA-A*03:01 complexes was determined through fluorescence anisotropy as previously described^45^. Briefly, purified peptide/HLA-A:03:01 complexes were generated using derivatives of the WT and Mut PI3Kα peptides where the P5 Gly position 5 was substituted with a 5-carboxyfluorescein-moodified lysine. Experiments were performed on a Beacon 2000 Fluorescence Polarization Instrument (Invitrogen). 100 nM fluorescein-labeled peptide/MHC complexes were mixed with 100 μM unlabeled peptide in 20 mM NaH_2_PO_4_ and 75 mM NaCl (pH 7.4). The excitation wavelength was 488 nm and anisotropy was detected at 535 nm. Changes in anisotropy were recorded as a function of time. Dissociation kinetics of peptides were determined by fitting the anisotropy curve to a single or biphasic exponential decay function in MATLAB. Half-lives were computed from the slowest dissociation rate from the relationship t_1/2_ = 0.693/k_off_.

### Protein crystallization

Purified proteins were exchanged into 10 mM HEPES and 20 mM NaCl (pH 7.4) before crystallization. Crystals of pWT/HLA-A*03:01 grew in 10% PEG 8000, 200 mM calcium acetate, 100 mM HEPES (pH 7.5) at a protein concentration of 15 mg/mL. Crystals of pMut/HLA-A*03:01 grew in 20% PEG 3350, 200 mM ammonium formate (pH 6.6) at a protein concentration of 15 mg mL^−1^. Crystals of the TCR4-pMut/HLA-A*03:01 complex grew in 10% PEG 8000, 200 mM magnesium acetate at a protein concentration of 5 mg mL^−1^. Crystals were obtained by hanging drop vapor diffusion at 4°C and were cryoprotected with 15% glycerol and flash-frozen before data collection.

### X-ray diffraction and structure determination

X-ray diffraction was performed at the beamline 22-ID or 24-ID-E of the Advanced Photon Source at Argonne National Laboratory. Diffraction data were processed with HKL2000 and solved by molecular replacement via Phaser in Phenix. The search model for the HLA-A*03:01 complexes was PDB 2XPG^81^. The search model for the complex with TCR4 was built by Sculptor with the α chain from PDB 3QH3 and the β chain from PDB 4PRH^82,83^. Peptides and for the TCR4 complex the CDR loops were removed before molecular replacement and manually rebuilt in Coot after the model was obtained from PHENIX Autobuild. Models were further refined automatically in PHENIX and manually in Coot. Structures were visualized using PyMOL 2.3.4 and Discovery Studio 2019.

### Surface plasmon resonance

Binding measurements between TCR4 and pWT or pMut/HLA-I complexes were performed using SPR on a Biacore T200 instrument as previously described^69^. Briefly, proteins were exchanged into 10 mM HEPES, 150 mM NaCl, 3 mM EDTA and 0.005% surfactant P-20, pH 7.4. TCR4 was immobilized on a CM5 Series S sensor chip to 900-1300 response units via amine coupling. Experiments were performed at 25°C with blank activated and deactivated flow cells as a reference. pMHC complexes were injected at a flow rate of 5 μl ml^−1^ as analytes in duplicate. Binding affinities were determined by fitting the steady-state data from both sets of injections to a 1:1 binding model in OriginPro.

### *PIK3CA* clonality, tumor phylogeny, and HLA LOH

The proportion of cancer cells harboring *PIK3CA* (H1047L) (cancer cell fraction, CCF) was estimated by integrating number of copies of the mutant allele, purity of the tumor and variant allele frequency of the mutation^84^. Allele-specific copy number inferences were made using FACETS^85^ and the number of copies of the mutant allele was inferred as described previously^86^. Mutations were deemed clonal if the CCF value is >0.8 or if the CCF value is >0.7 and the upper bound of the 95^th^-percentile confidence interval is >0.9. All mutations for which CCF was determinable but that did not meet the criteria for being clonal were designated as sub-clonal. HLA genotypes from MSK-IMPACT were inferred using POLYSOLVER^56,87^. Loss of heterozygosity (LOH) at HLA Class I alleles was inferred using LOHHLA^62^. At each of the HLA Class I loci, LOH was called if the median copy number calculated by LOHHLA is less than 0.5 and the *P*-value reflecting the allelic imbalance was less than 0.001.

## ACKNOWLEDGEMENTS

This study was supported in part by NIH R37 CA259177-01 (C.A.K.), NIH P30 CA008748 (C.A.K. and D.A.S.), the Damon Runyon Cancer Research Foundation CI-96-18 (C.A.K.), Cancer Research Institute CRI3176 (C.A.K), Functional Genomics Initiative Rapid Response Pilot Grant (C.A.K), The Manhasset Women’s Coalition Against Breast Cancer (C.A.K.), the Emerson Collective Cancer Fund (C.A.K.), and a sponsored research agreement with Intima Bioscience (C.A.K.). X-ray diffraction data were collected at the Advanced Photon Source, supported by DOE contract DE-AC02-06CH11357, and the NE-CAT and SER-CAT beamlines, supported by member institutions and NIH grants P30GM124165, S10OD021527, S10RR25528, and S10RR028976. We thank Alex Kentsis for access to the Byonic software and the Proteomics Resource Center at Rockefeller University for the performance of all LC-MS/MS experiments. We thank the Epigenomics Core of Weill Cornell Medical College for enabling single-cell immunoprofiling experiments. We thank Matthew R. Femia for technical assistance with pilot IVS screens.

## Author contributions

S.S.C. performed all *in vitro* sensitization, TCR cloning, TCR transduction, and TCR functional assays with technical assistance from S.N.F. Single-cell sequencing analysis was performed by F.D., P.Z. and D.B. IP/LC-MS/MS experiments were performed and analyzed by M.G.K. and D.A.S. Crystallographic, S.P.R., p/HLA-I thermal, and p/HLA-I kinetic stability experiments were performed by J.M. and B.M.B. Determination of CCF were performed by C.B. and P.R; HLA-I calls, HLA-LOH, antigen processing and presentation mutations, and phylogenetic analyses were performed by C.B. Pathology review and tissues processing was performed H.Y.M., B.W., and P.R.; H.Y.M. performed TIL calls. *PIK3CA* public NeoAg biospecimen identification, collection, and archiving was performed by L.B.B., C.B., S.S.C., S.N.F., W.D.B., and A.D. The manuscript was written by S.S.C., B.M.B, and C.A.K. with editorial input from all co-authors. S.S.C. and C.A.K. supervised and coordinated all aspects of this work.

## Data availability

All data generated and supporting the findings of this study are available within the paper. Single-cell RNA-seq datasets will be deposited in the Gene Expression Omnibus (GEO) prior to publication. Structural data, including coordinates and structure factors, of pWT/HLA-I, pMut/HLA-I and the ternary TCR4/pMut/HLA-I complex are available at the Protein Data Bank (https://www.rcsb.org/) under PDB accession codes 7L1B, 7L1C and 7L1D. Additional information and materials will be made available upon request.

## Code availability

Custom R scripts to analyze single-cell RNA-seq and V(D)J Immunoprofiling data sets will be deposited in https://github.com/abcwcm and made available upon request.

## EXTENDED DATA FIGURE LEGENDS

**Extended Data Fig. 1: Negative and positive predictive capacity of the SIFT-seq discovery platform for Mut *PIK3CA*-specific TCRs.** Volcano plots displaying global transcriptomic changes for (**A**) MSK 21LT2 clonotype 18 (non-reactive; frequency rank = 1), (**B**) MSK 0606T clonotype 367 (reactive; frequency rank = 23), and (**C**) MSK 0606T clonotype 115 (non-reactive; frequency rank = 1) following stimulation with Mut versus WT *PIK3CA*. Vertical and horizontal dashed lines indicate thresholds for gene expression fold change (FC) and statistical significance, respectively. Orange and blue dots represent significantly up and down-regulated genes following Mut *PIK3CA* stimulation, respectively. (**D**) Representative FACS plots and (**E**) summary bar graph displaying the frequency of intra-cellular IL-2 production in open-repertoire T cells retrovirally transduced with MSK 21LT2 clonotype 18. Transduced T cells (live^+^mTCR^+^CD3^+^) were co-cultured with autologous monocyte derived DCs electroporated with mRNA encoding Mut or WT *PIK3CA*. Stimulation by phorbol 12-myristate 13-acetate-ionomycin (PMA-Iono) was used as a TCR-independent positive control for T cell function. Bar graphs displayed as mean ± SEM using *n*=3 replicates per condition. *** = *P* ≤ 0.001. ns = not significant. Two-sided Student’s t-test with Bonferroni correction.

**Extended Data Fig. 2: Variable chain gene sequences and HLA specificity of a *PIK3CA* public NeoAg-specific TCR library.** (**A**) Table listing *TRAV*, *TRAJ*, *TRBV* and *TRBJ* gene segment composition and CDR3α/β lengths for a library of *PIK3CA* public NeoAg-specific TCRs retrieved using the SIFT-seq discovery platform. (**B**) Representative FACS plots and (**C**) summary bar graphs demonstrating the specificity of *PIK3CA* public NeoAg TCRs 1-4 for HLA-A03 supertype members HLA-A*03:01, -A*03:02, and -A*11:01. The number of shared amino acids and % homology of each supertype member to HLA-A*03:01 are shown. T cells transduced with individual TCR library members were cocultured with mono-allelic cell lines co-expressing the indicated HLA-A03 supertype member and either Mut or WT *PIK3CA*. Numbers within each FACS plot and bar graph indicate the frequency of TNFα producing T cells after pre-gating on live^+^mTCR^+^CD8^+^ cells. Bar graphs displayed as mean ± SEM using *n*=3 replicates per condition. Data shown is presentative of 2 independent experiments.

**Extended Data Fig. 3: Comparison of mutation-specific effector functions for a library of *PIK3CA* public NeoAg-specific TCRs.** (**A**) Summary bar graphs demonstrating Mut-specific expression of multiple effector molecules by T cells transduced with a *PIK3CA* public NeoAg TCR. Open-repertoire CD8^+^ T cells were individually transduced with *PIK3CA* public NeoAg TCR library members 1-4 and cocultured with HLA-A*03:01^+^ target cells that express either WT or Mut *PIK3CA*. Numbers in each bar graph indicate the frequency of T cells producing TNFα, IFNγ, or CD107a measured by FACS analysis after pre-gating on live^+^mTCR^+^CD8^+^ cells. Bar graphs are displayed as mean ± SEM using *n*=3 replicates per condition. (**B**) Mean effective concentration of Mut *PIK3CA* required to trigger half-maximal (EC_50_) TNFα production in enriched CD8^+^ or CD4^+^ T cells transduced with a *PIK3CA* public NeoAg TCR. CD8^+^ or CD4^+^ T cells were individually transduced with *PIK3CA* public NeoAg TCRs 1-4. Transduced T cells were co-cultured with an HLA-I mono-allelic cell line expressing HLA-A*03:01 and electroporated with defined concentrations of Mut *PIK3CA* mRNA. All data shown is representative of 2 independent experiments.

**Extended Data Fig. 4: Structural details and comparisons of the PI3Kα pMut and pWT peptides bound to HLA-A*03:01.** (**A** and **B**) Electron density of the pMut and pWT peptides in the HLA-A:03:01 (A*03:01) binding groove with 2F_o_-F_c_ contoured at 1σ. (**C**) Comparison of the peptide conformations in the binding grooves. A small structural divergence is present at the backbone at P4 and P5 Gly residues but does not alter overall peptide conformation or amino acid side chains positions. (**D**) Conceptual model of the peptides modified at P5 with fluorescein derivatized lysines (K-Flc). The fluorescent group is not expected to interfere with the interaction of the peptide with the HLA-I molecule.

**Extended Data Fig. 5: Structural details of the TCR4/pMut/HLA-A*03:01 complex.** (**A**) Electron density of the interface with 2F_o_-F_c_ electron density contoured at 1σ for the peptide and CDR3α/CDR3β loops. (**B**) Additional views emphasizing the position and architecture of CDR3β relative to the pMut/HLA-I complex. (**C**) View of the conformational change that occurs in the pMut peptide upon TCR4 binding, centered on the P6 Trp side chain. The change occurs predominantly through a rotation of the pG5 ϕ angle from 75° to −167° upon TCR4 binding.

**Extended Data Fig. 6: Application of MSK IMPACT clinical next-gen sequencing (NGS) for the efficient identification of *PIK3CA* public NeoAg-expressing cancer patients.** (**A**) MSK IMPACT is a CLIA-certified clinical NGS platform capable of performing high resolution HLA-I calls. Using DARWIN, an automated genotype enrollment tool, patients who have a *PIK3CA* (H1047L) (Mut^+^) cancer and are HLA-A*03:01 (HLA^+^) are flagged for biospecimen collection using an IRB approved protocol. (**B**) Using this approach, a biotrust of viable peripheral blood mononuclear cells (PBMCs) from a cohort of cancer patients with diverse malignancies who express an identical *PIK3CA* public neoantigen (NeoAg) was collected. (**C**) Summary table of *PIK3CA* public NeoAg expressing cancer types analyzed in this cohort.

