## Supplemental figures and tables for "Immunogenicity of a public neoantigen derived from mutated *PIK3CA*"

**Extended Data Fig. 1:** Negative and positive predictive capacity of the SIFT-seq discovery platform for Mut *PIK3CA*-specific TCRs.

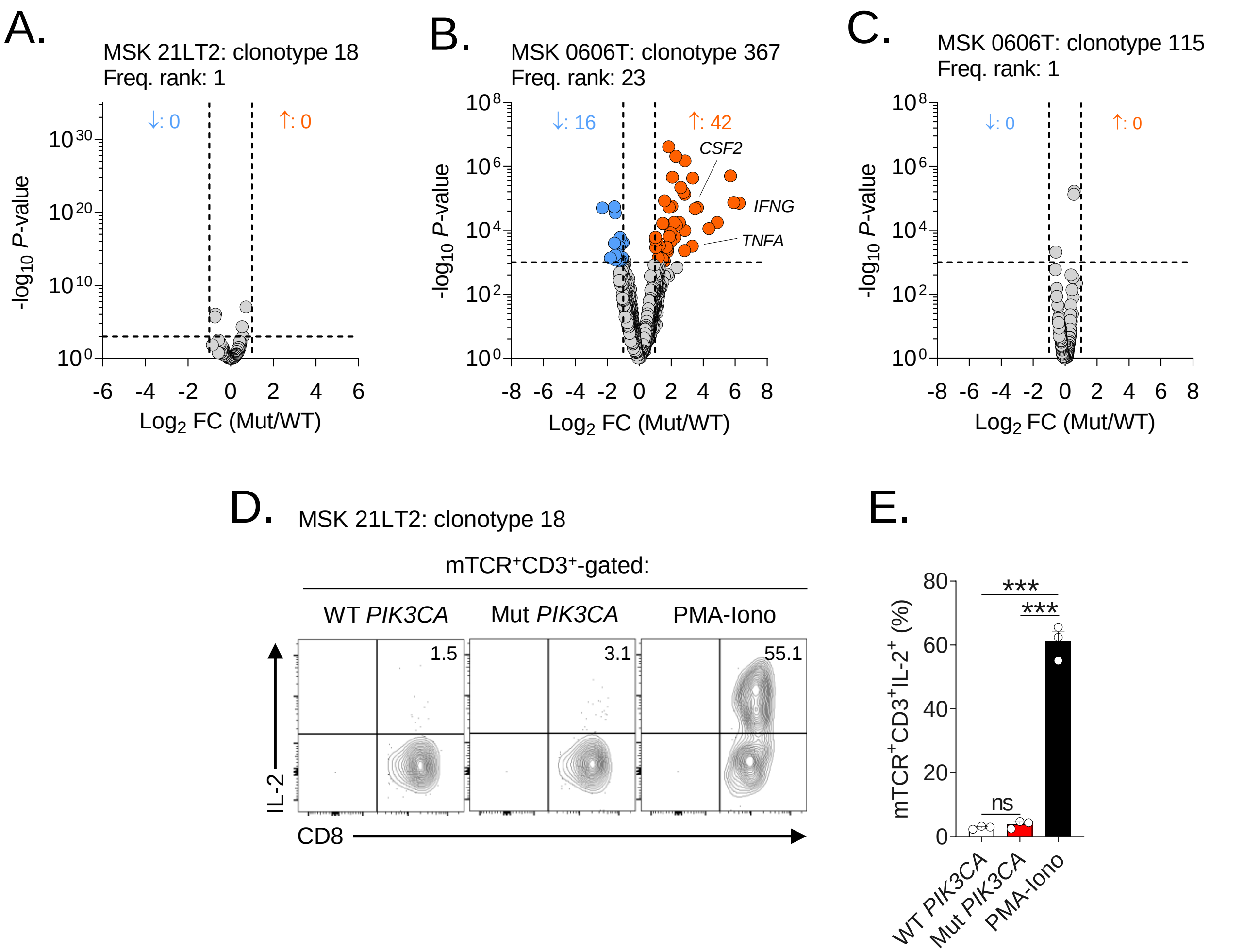

**Extended Data Fig. 2:** Variable chain gene sequences and HLA specificity of a *PIK3CA* public NeoAg-specific TCR library.

A.

| TCR ID | TRAV | TRAJ | TRBV | TRBJ | CDR3α length | CDR3β length |
|---|---|---|---|---|---|---|
| TCR1 | TRAV26-1 | TRAJ22 | TRBV5-6 | TRBJ2-7 | 14 | 14 |
| TCR2 | TRAV9-2 | TRAJ6 | TRBV4-1 | TRBJ2-5 | 13 | 16 |
| TCR3 | TRAV4 | TRAJ9 | TRBV11-2 | TRBJ2-7 | 15 | 14 |
| TCR4 | TRAV12-2 | TRAJ44 | TRBV9 | TRBJ2-6 | 13 | 19 |

B.

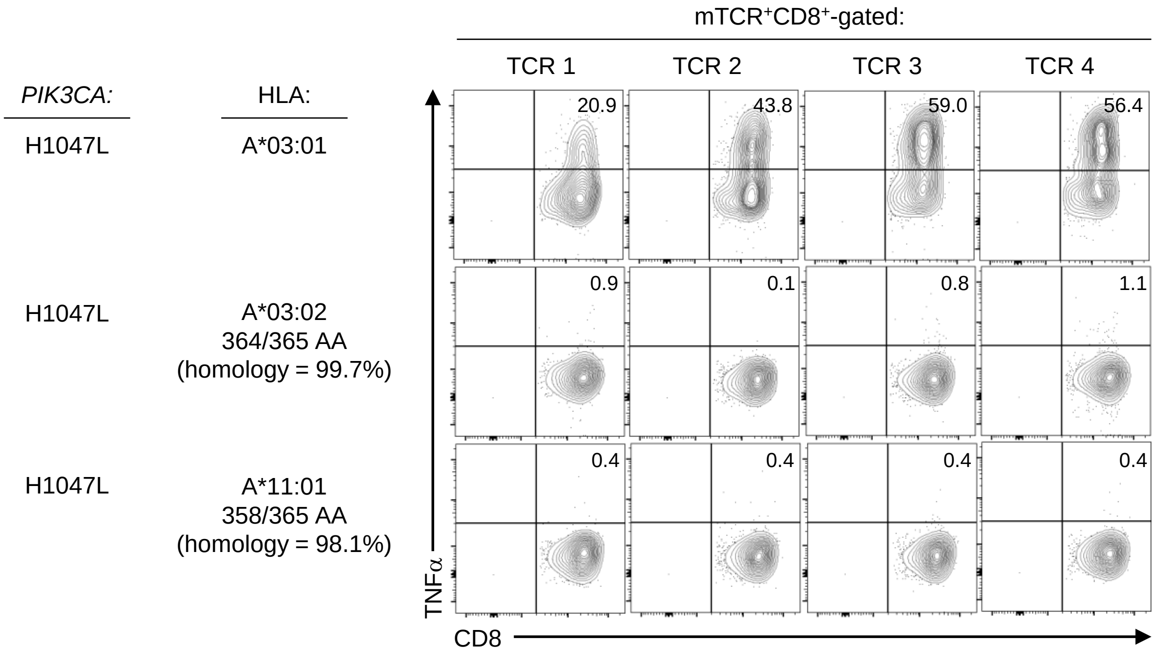


C.

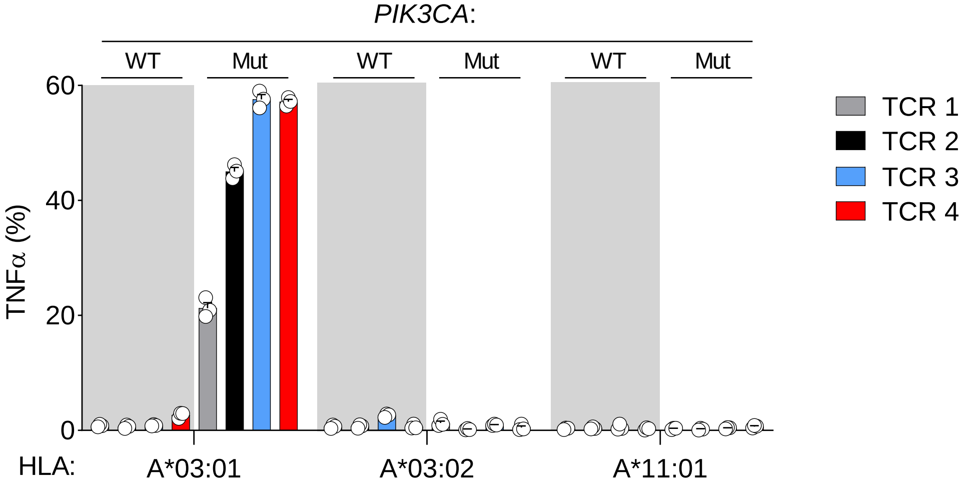

**Extended Data Fig. 3:** Comparison of mutation-specific effector functions for a library of *PIK3CA* public neoantigen-specific TCRs.

A.

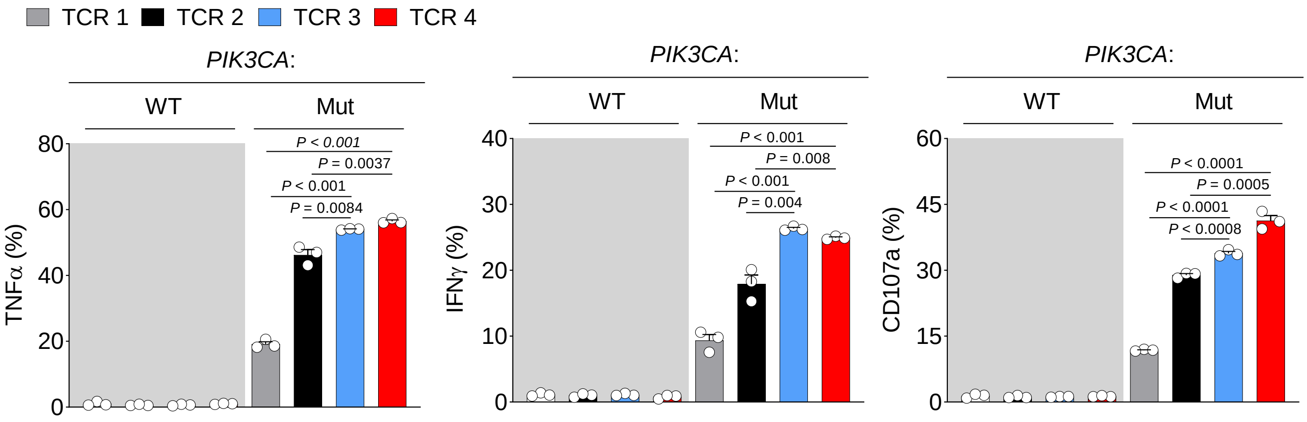


B.

| TCR ID | $EC_{50}$ $CD8^+$ (μM) | $EC_{50}$ $CD4^+$ (μM) |
|---|---|---|
| TCR1 | 169 | - |
| TCR2 | 0.268 | - |
| TCR3 | 0.029 | 0.489 |
| TCR4 | 0.026 | 0.769 |

**Supplementary Table 1:** X-ray data collection and refinement statistics for the pWT and pMut peptide/HLA-A*03:01 (A*03:01) complexes and the complex of pMut/A*03:01 with TCR4 crystal structures.

| | **pWT/A*03:01** | **pMut/A*03:01** | **TCR4-pMut/A*03:01** |
|---|---|---|---|
| **Data Collection** | | | |
| Space group | P 6 2 2 | P 6 2 2 | P 43 21 2 |
| Unit cell dimensions | | | |
| *a, b, c* (Å) | 156.59, 156.59, 86.21 | 155.72, 155.72, 85.66 | 72.28, 72.28, 475.73 |
| *α, β, γ* (°) | 90, 90, 120 | 90, 90, 120 | 90, 90, 90 |
| Resolution (Å) | 50.00-2.05 (2.09-2.05) | 50.00-1.96 (1.99-1.96) | 50.00-3.11 (3.15-3.11) |
| $R_{merge}$ | 0.110 (0.598) | 0.123 (0.918) | 0.145 (1.365) |
| $I/\sigma I$ | 33.7 (2.5) | 29.3 (2.6) | 25.0 (2.0) |
| Completeness (%) | 98.9 (89.0) | 100 (100) | 99.9 (97.6) |
| Total reflections | 1,579,824 | 1,911,199 | 1,856,431 |
| Unique Reflections | 39,401 (1745) | 44,346 (2187) | 24,158 (914) |
| Redundancy | 17.5 (11.7) | 17.6 (10.9) | 20.2 (14.1) |
| **Refinement** | | | |
| Resolution (Å) | 45.20-2.04 | 44.95-1.96 | 49.51-3.11 |
| $R_{work}/R_{free}$ | 0.195/0.221 | 0.169/0.205 | 0.198/0.250 |
| No. atoms | | | |
| Protein | 3430 | 3484 | 6512 |
| Average B-factors (Å) | 45.0 | 37.0 | 98.0 |
| r.m.s. deviations | | | |
| Bond length (Å) | 0.002 | 0.015 | 0.002 |
| Bond angles (°) | 0.457 | 1.189 | 0.494 |
| Ramachandran favored (%) | 98.14 | 98.94 | 97.64 |
| Ramachandran outliers (%) | 0 | 0 | 0 |
| PDB accession code | 7L1B | 7L1C | 7L1D |

**Extended Data Fig. 4:** Structural details and comparisons of the PI3Kα pMut and pWT peptides bound to HLA-A*03:01.

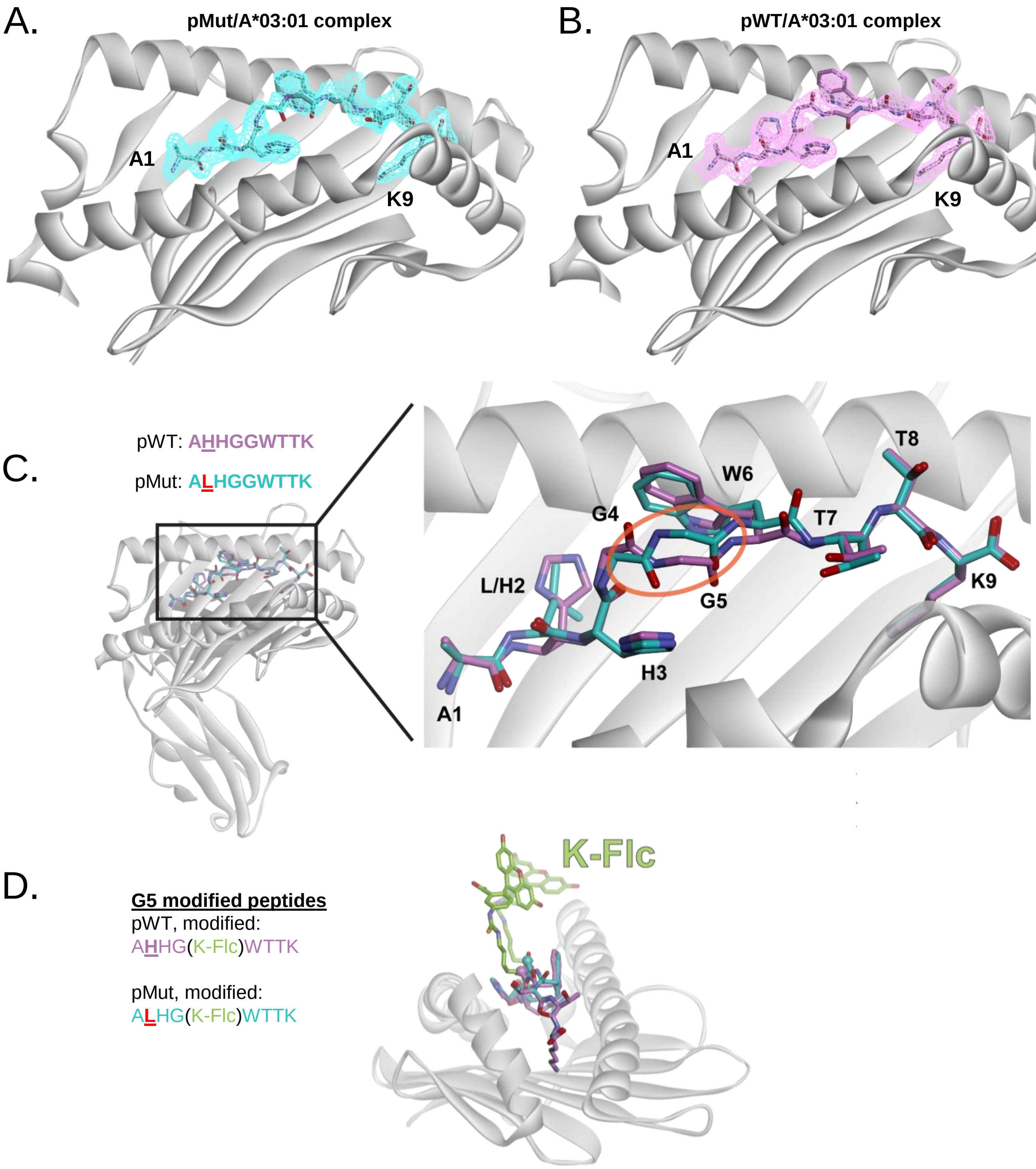

**Extended Data Fig. 5:** Structural details of the TCR4/pMut/HLA-A*03:01 complex.

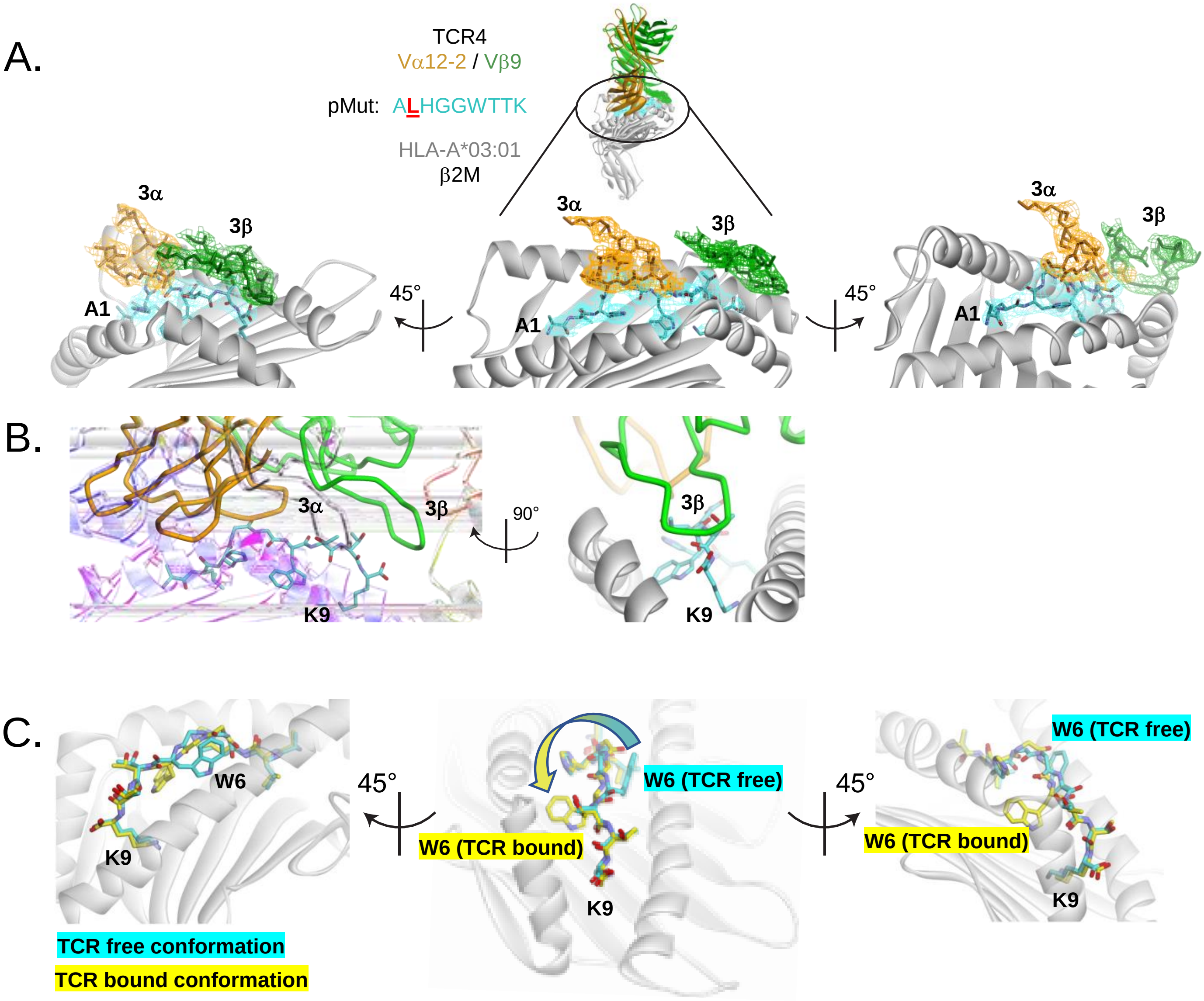

**Supplementary Table 2:** Assessment of the cross-reactivity potential for TCR4.

| Gene | Taxonomy ID | Length | Position | Sequence |
|---|---|---|---|---|
| *PIK3CA* (H1047L) | n.a. | 1068 | 1046-1054 | A**L**HG<u>GWTTK</u> |
| WT *PIK3CA* | P42336 | 1068 | 1046-1054 | A**H**HG<u>GWTTK</u> |
| *TM87B* | Q96K49 | 555 | 407-415 | IVFM<u>GWTTK</u> |

Derived from ScanProsite using the motif ‘x-x-x-x-G-W-T-T-K’
n.a. = not applicable

**Extended Data Fig. 6:** Application of MSK IMPACT clinical next-gen sequencing (NGS) for the efficient identification of *PIK3CA* public NeoAg-expressing cancer patients.

A.

**MSK IMPACT clinical NGS**

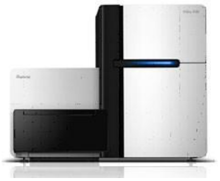

- High-res HLA-I typing
- Weekly email alerts for Mut+/HLA+ patients (**DARWIN**)

B.

*PIK3CA* (H1047L)+ / HLA-A*03:01+ cancer patients

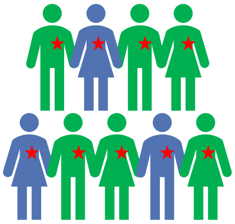

*PIK3CA* public neoantigen cohort, *n*=14

PBMC collection

C.

| HLA-A | *PIK3CA* | Disease | (*n*) |
|---|---|---|---|
| *03:01 | H1047L | BRCA | 8 |
| | H1047L | UCEC | 1 |
| | H1047L | COAD/ READ | 2 |
| | H1047L | Other | 3 |

**Supplementary Table 3:** Demographics of HLA-A*03:01$^{+}$ cancer patients expressing the *PIK3CA* (H1047L) public neoantigen.

| Subject ID | *PIK3CA* | HLA-A, allele 1 | HLA-A, allele 2 | Cancer type | Disease status | TIL | Reactive | Prior treatment(s) |
|---|---|---|---|---|---|---|---|---|
| 1 | H1047L | *03:01 | *02:01 | BRCA | Met | No | No | chemo, ET, IO, PIK3CA(i), XRT |
| 2 | H1047L | *03:01 | *32:01 | BRCA | NED | Yes | Yes | chemo, ET, XRT |
| 3 | H1047L | *03:01 | *26:01 | BRCA | NED | Yes | No | ET |
| 4 | H1047L | *03:01 | *68:01 | BRCA | NED | Yes | Yes | chemo, ET, XRT |
| 5 | H1047L | *03:01 | *02:01 | BRCA | Met | No | No | chemo, ET, PIK3CA(i) |
| 6 | H1047L | *03:01 | *23:01 | BRCA | NED | Yes | No | ET |
| 7 | H1047L | *03:01 | *32:01 | BRCA | NED | Yes | No | ET, XRT |
| 8 | H1047L | *03:01 | *02:02 | BRCA | Met | No | No | ET, anti-HER2 |
| 9 | H1047L | *03:01 | *02:01 | UCEC | Met | Yes | No | chemo, IO |
| 10 | H1047L | *03:01 | *03:01 | COAD | Met | No | No | chemo, anti-VEGF |
| 11 | H1047L | *03:01 | *02:01 | COAD | Met | Yes | No | chemo |
| 12 | H1047L | *03:01 | *23:01 | GBM | Recurrent | No | No | chemo, XRT |
| 13 | H1047L | *03:01 | *02:06 | THCA | Met | No | Yes | $I^{131}$, IO, TKI |
| 14 | H1047L | *03:01 | *33:01 | BLCA | NED | No | Yes | None |

BRAC = breast cancer; UCEC = uterine corpus endometrial cancer; COAD = colon adenocarcinoma;
GBM = glioblastoma multiforme; THCA = thyroid cancer; BLCA = bladder cancer.
chemo = chemotherapy; ET = endocrine therapy; IO = immunotherapy; PIK3CA(i) = PIK3CA inhibitor;
XRT = radiation therapy; TKI = tyrosine kinase inhibitor; $I^{131}$ = radioactive iodine.
Met = metastatic; NED = no evaluable disease.

**Extended Data Fig.7:** Representative gating strategy to determine NeoAg-specific cytokine production of TCR-transduced T cells

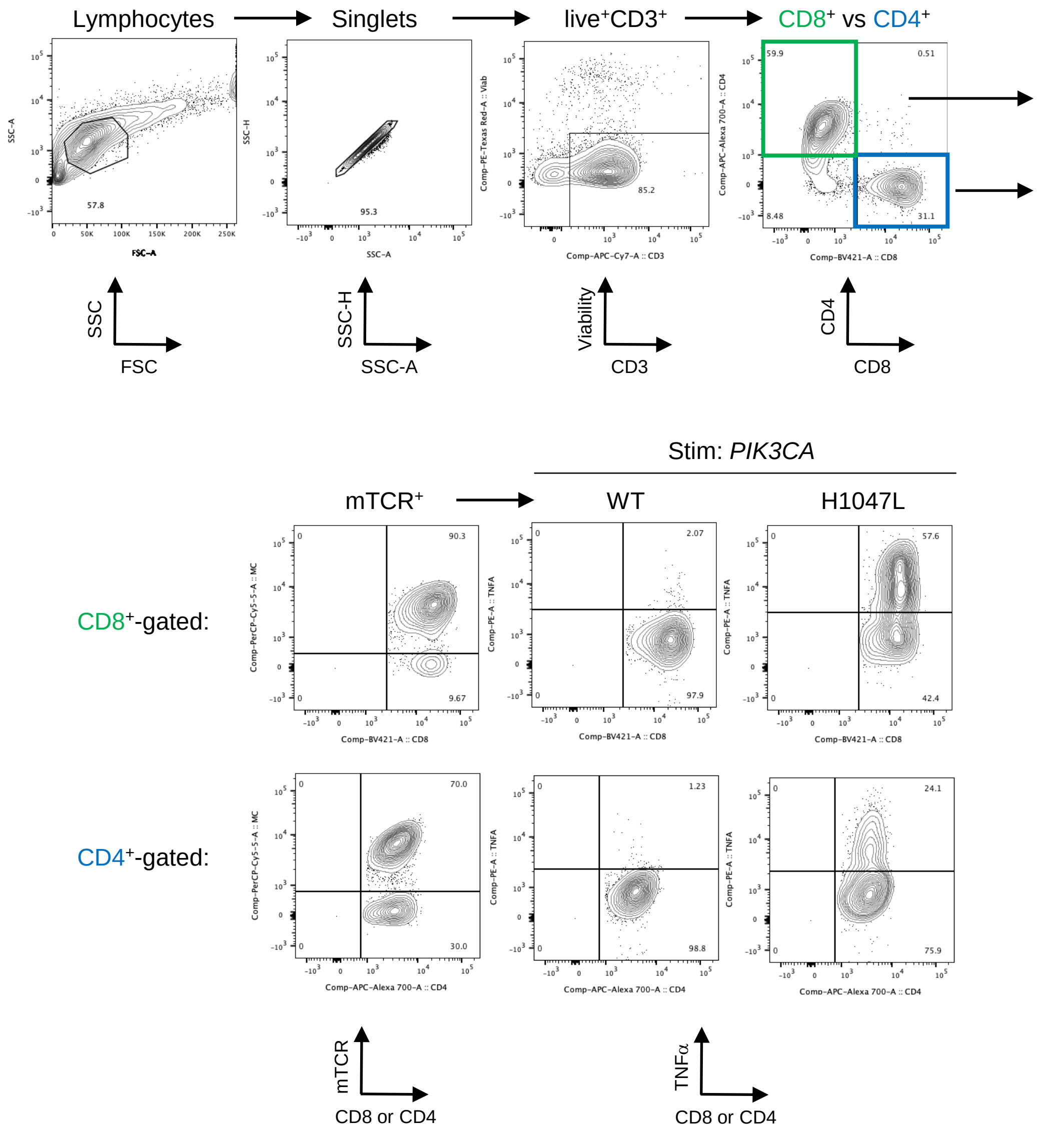
